# Characterization of the brain functional architecture of psychostimulant withdrawal using single-cell whole brain imaging

**DOI:** 10.1101/743799

**Authors:** Adam Kimbrough, Lauren C. Smith, Marsida Kallupi, Sierra Simpson, Andres Collazo, Olivier George

**Affiliations:** School of Medicine, Department of Psychiatry, University of California San Diego, MC 0667, La Jolla, California, 92093; Department of Neuroscience, The Scripps Research Institute, La Jolla, California 92037; Beckman Institute, Cal-Tech, MC 139-74, Pasadena, California, 91125

## Abstract

Numerous brain regions have been identified as contributing to addiction-like behaviors, but unclear is the way in which these brain regions as a whole lead to addiction. The search for a final common brain pathway that is involved in addiction remains elusive. To address this question, we used male C57BL/6J mice and performed single-cell whole-brain imaging of neural activity during withdrawal from cocaine, methamphetamine, and nicotine. We used hierarchical clustering and graph theory to identify similarities and differences in brain functional architecture. Although methamphetamine and cocaine shared some network similarities, the main common neuroadaptation between these psychostimulant drugs was a dramatic decrease in modularity, with a shift from a cortical- to subcortical-driven network, including a decrease in total hub brain regions. These results demonstrate that psychostimulant withdrawal produces the drug-dependent remodeling of functional architecture of the brain and suggest that the decreased modularity of brain functional networks and not a specific set of brain regions may represent the final common pathway that leads to addiction.

**Significance Statement:** A key aspect of treating drug abuse is understanding similarities and differences of how drugs of abuse affect the brain. In the present study we examined how the brain is altered during withdrawal from psychostimulants. We found that each drug produced a unique pattern of activity in the brain, but that brains in withdrawal from cocaine and methamphetamine shared similar features. Interestingly, we found the major common link between withdrawal from all psychostimulants, when compared to controls, was a shift in the broad organization of the brain in the form of reduced modularity. Reduced modularity has been shown in several brain disorders, including traumatic brain injury, and dementia, and may be the common link between drugs of abuse.

## Introduction

Psychostimulants are a class of highly addictive and commonly abused drugs that includes cocaine, nicotine, and methamphetamine [1, 2]. A large number of brain regions have been implicated in dependence and addiction-like behaviors that are associated with psychostimulant use [3–9]. However, the complete neural network that is associated with psychostimulant withdrawal remains understudied, and the search for a common brain pathway that is responsible for psychostimulant withdrawal remains elusive. Common features of dependence may not be found at the brain region level but rather at the network level.

The identification of changes in neural network structure that are caused by psychostimulant withdrawal may be critical to understanding the ways in which these drugs affect the brain in a common way. Previous studies identified changes in network function after psychostimulant use [10–13], but these analyses focused on macroscale changes and not the mesoscale level, or they focused on preselected regions of interest.

The present study sought to identify the ways in which withdrawal from different commonly abused psychostimulants alters functional architecture of the brain. We hypothesized that withdrawal from psychostimulants would result in major changes in functional neural networks and decrease modular structuring of the brain. We further hypothesized that each psychostimulant that was examined herein (i.e., methamphetamine, nicotine, and cocaine) would have a unique neural network that is associated with withdrawal. We measured single-cell whole-brain activity using Fos as a marker for neuronal activation in mice that underwent withdrawal from chronic psychostimulant (cocaine, methamphetamine, and nicotine) administration. The psychostimulant doses were chosen based on previous studies that reported rewarding effects during use and observed withdrawal-like symptoms after the cessation of chronic exposure for each drug [14–20]. We then used single-cell whole-brain activity to identify coactivation patterns of brain regions in the network that was associated with each treatment using hierarchical clustering. The coactivation patterns were used to determine the modular structuring of each network. Graph theory was then used to further characterize each network to determine the brain regions that are responsible for intra- and intermodular connectivity.

## Results

### Psychostimulant withdrawal induces massive restructuring of the brain

We examined the ways in which withdrawal from different psychostimulants alters neural coactivation and modular structuring of the brain. For an overview of the experimental design, see Fig. 1A. For all of the drugs tested, acute withdrawal produced widespread increases in the coactivation of brain regions compared with saline controls (Fig. 1C-F). Importantly, modular structuring of the brain decreased in response to withdrawal from each psychostimulant compared with controls. When using a threshold of 50% of tree height, saline control mice exhibited a modular structure of the brain that contained seven modules, whereas cocaine mice had four modules, methamphetamine mice had three modules, and nicotine mice had five modules and one isolated brain region that was not grouped with any other region (i.e., interanterodorsal nucleus of the thalamus; Fig. 1B-F). Notably, the decrease in the number of modules during withdrawal was independent of the clustering thresholds that were used (Fig. 1B). These data indicate that psychostimulant withdrawal decreases modularity of the brain functional network compared with controls.

**Figure 1.**
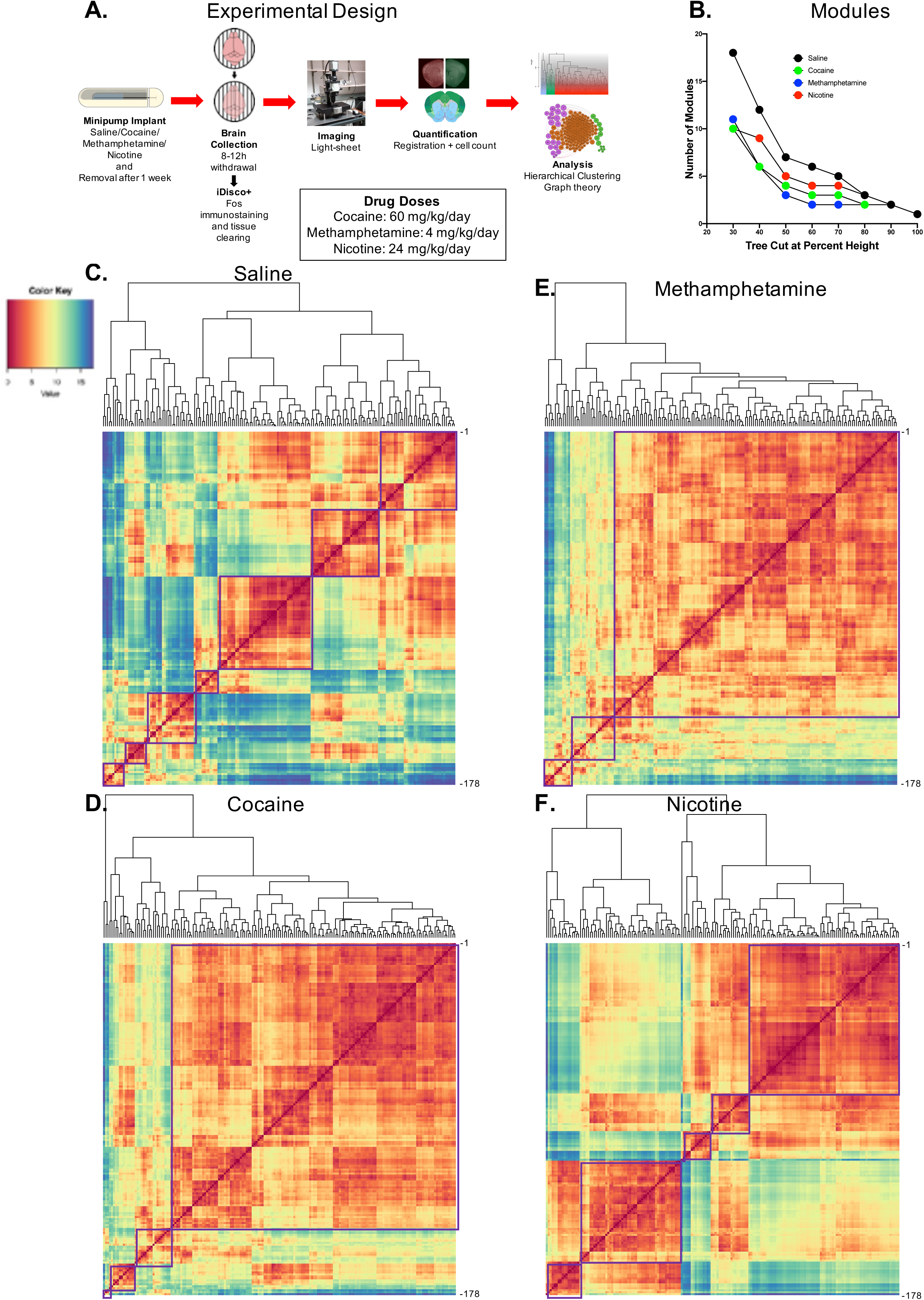
**A.** Experimental design. Mice were surgically implanted with an osmotic minipump that contained either saline or a psychostimulant (60 mg/kg/day cocaine, 4 mg/kg/day methamphetamine, or 24 mg/kg/day nicotine). They were then returned to their home cage for 1 week. After 1 week, the minipumps were surgically removed, and the mice were returned to their home cage until brain tissue was collected 8 h later (saline, cocaine, nicotine) or 12 h later (methamphetamine). Brains were then processed for whole-brain Fos immunohistochemistry and clearing via iDISCO+ and then imaged on a light-sheet microscope. Fos values were detected and registered to the Allen Brain Atlas using ClearMap [63]. Pearson correlations were then calculated to determine functional coactivation among brain regions. Brain regions were then grouped into modules based on their coactivation patterns through hierarchical clustering. Graph theory analyses was then performed to identify brain regions that are heavily involved in intra- and intermodular connectivity. **B-F.** Hierarchical clustering of complete Euclidean distance matrices for each treatment. Modules were determined by cutting each dendrogram at half of the maximal tree height. **B.** Number of modules in each treatment condition after cutting the hierarchical clustered dendrogram at different percentages of tree height. In all cases (except at extreme cutoff values; e.g., 90-100%), the psychostimulant networks showed lower modularity compared with the control network. **C.** Relative distance of each brain region relative to the others that were examined in control mice. In control mice, seven distinct modules of coactivation were identified. **D.** Relative distance of each brain region relative to the others that were examined in cocaine mice. In cocaine mice, four distinct modules of coactivation were identified. **E.** Relative distance of each brain region relative to the others that were examined in methamphetamine mice. In methamphetamine mice, three distinct modules of coactivation were identified. **F.** Relative distance of each brain region relative to the others that were examined in nicotine mice. In nicotine mice, five distinct modules of coactivation were identified. For all distance matrices, each module is boxed in purple. For the individual brain regions that are listed in panels C-F, see Table 6.

### Characterization of individual network features

To further characterize the features of each individual network, we used a graph theory approach to identify potential hub brain regions with the most intramodular and intermodular connectivity, which may drive activity within the network and thus be critical for neuronal function in the withdrawal state. We examined positive connectivity (thresholded to a Pearson correlation coefficient > 0.75 [0.75R] for inclusion as a network connection) for the network for each treatment and used the modular organization that was identified by hierarchical clustering to partition the regions of the networks. The 0.75R threshold was chosen because all of the brain regions in each network showed connections to other regions at this threshold. Previous animal model studies used various thresholds, ranging from 0.3R to 0.85R [21, 22], to examine connectivity. Negative network connectivity was not examined herein because the precise meaning of such connectivity is controversial and thus is not often examined in network-based approaches [23–26].

We determined the participation coefficient (PC; i.e., a measure of importance for intermodular connectivity) and the within-module degree Z-score (WMDz; i.e., a measure of importance for intramodular connectivity) [27] for all brain regions in the networks. A high PC was considered ≥ 0.30, and a high WMDz was considered ≥ 0.80. Overall, the control and nicotine networks showed much greater intermodular connectivity (high PC) and a great number of regions with both high intermodular and intramodular connectivity (high PC and WMDz). The cocaine and methamphetamine networks showed higher levels of intramodular connectivity (high WMDz) and a low number of regions with intermodular connectivity (Fig. 2A-C). We named each module in each network based on the group of brain regions with the highest WMDz score in the module and considered these regions to be drivers of activity within individual modules (see Fig. 3-6 for names).

**Figure 2.**
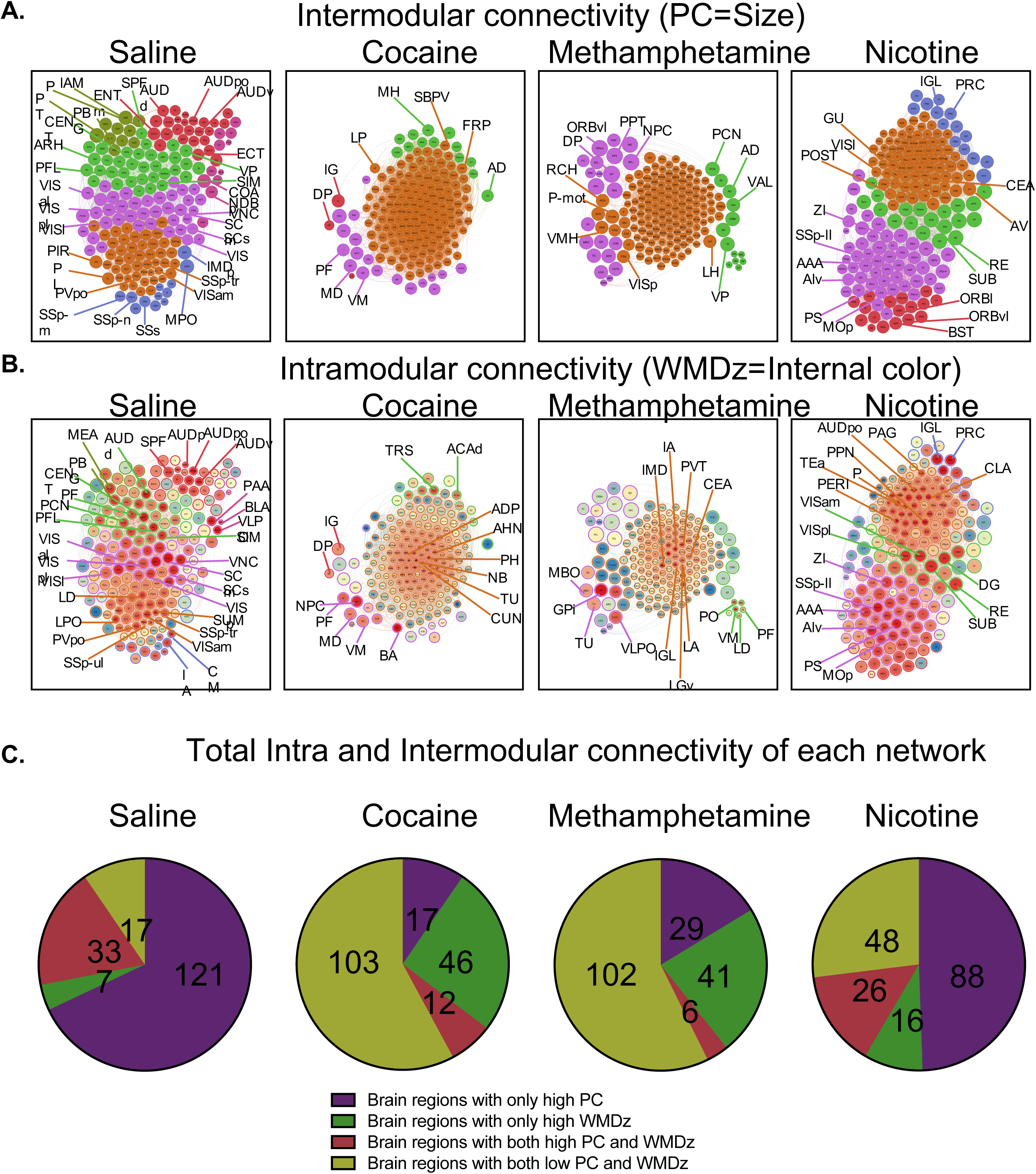
Intramodular (WMDz) and intermodular (PC) network features of each treatment. A high PC was considered ≥ 0.30, and a high WMDz was considered ≥ 0.80. **A.** Highlights of several regions with high PC in each module of each network network (see Table 1 for names of abbreviations). **B.** Highlights of several regions with high WMDz (red = higher, blue = lower) in each module of each network. Note that the WMDz color intensity is only relative to the other regions within the same network and not other networks (see Table 1 for names of abbreviations). **C.** Total number of brain regions that accounted for high PC, high WMDz, or both in each network. The control and nicotine networks showed much greater intermodular connectivity and a greater number of regions with both high intermodular and intramodular connectivity. The cocaine and methamphetamine networks showed higher levels of intramodular connectivity and a low number of regions with intermodular connectivity.

**Figure 3.**
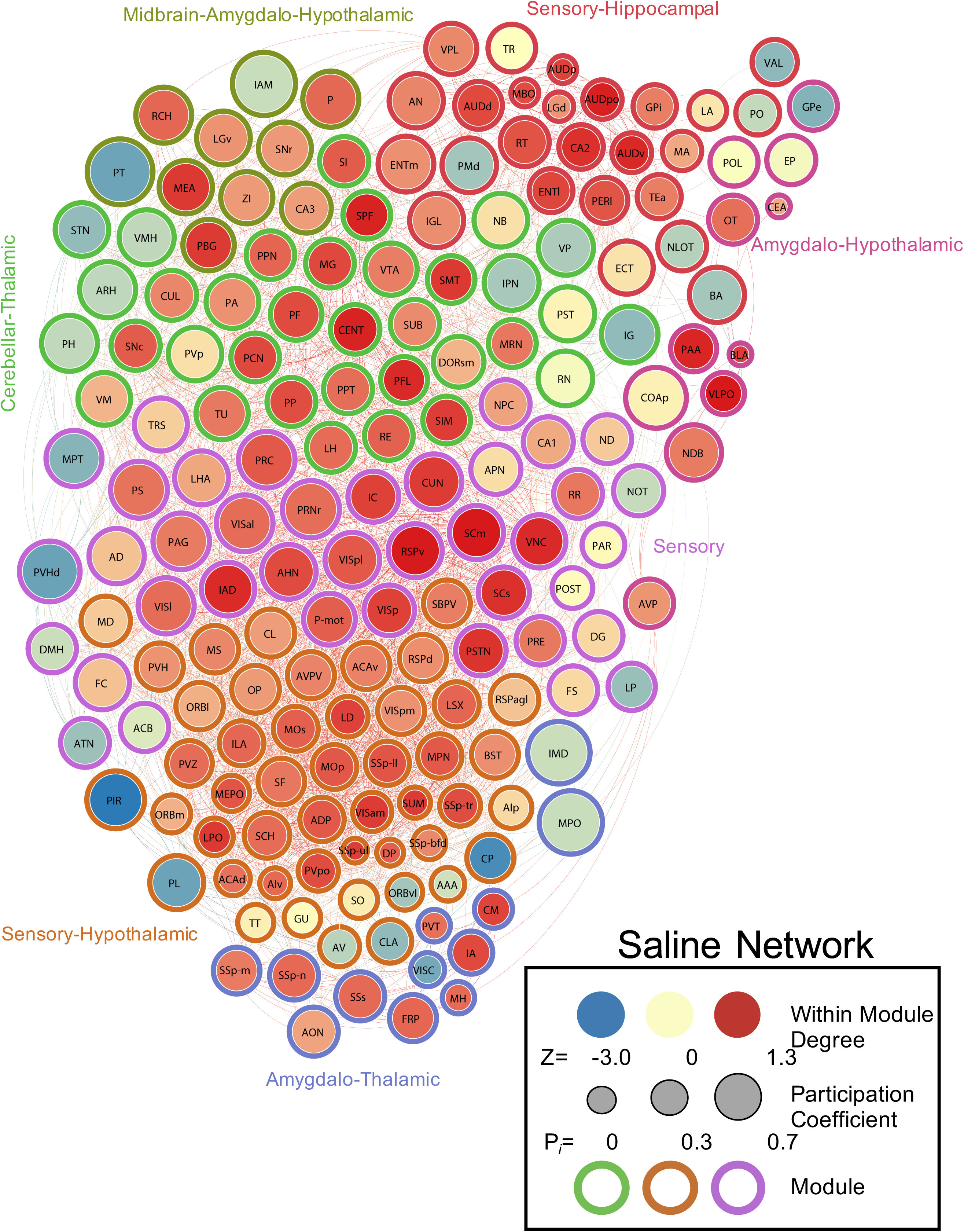
Neural network of control mice thresholded to 0.75R. Nodes/brain regions of the network are represented by circles. The size of the node represents the participation coefficient (smaller = lower PC; larger = higher PC). The internal color of each circle represents the within-module degree Z-score (dark blue = lowest; dark red = highest). The color of the modules that are identified in Fig. 1C are represented by different colored edges. See figure key for examples of each representative component of the figure.

**Figure 6.**
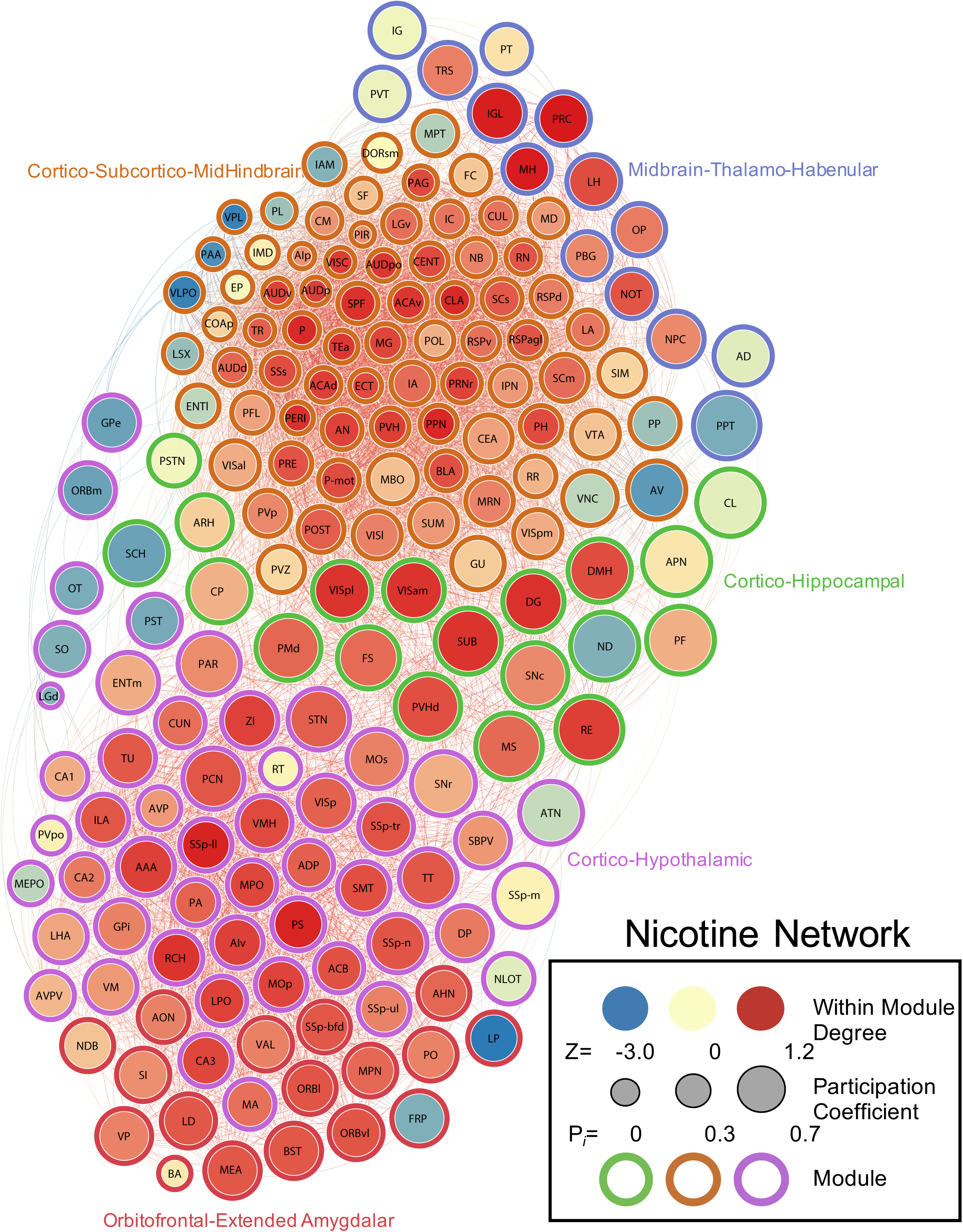
Neural network of nicotine mice during withdrawal thresholded to 0.75R. Nodes/brain regions of the network are represented by circles. The size of the node represents the participation coefficient (smaller = lower PC; larger = higher PC). The internal color of each circle represents the within-module degree Z-score (dark blue = lowest; dark red = highest). The color of the modules that are identified in Fig. 1F are represented by different colored edges. See figure key for examples of each representative component of the figure.

### The control network is driven by sensory-motor regions

The saline control network had 3,176 total connections and consisted of seven modules, many of which were heavily driven by sensory-motor brain regions. Of these seven modules, five contained several sensory or motor brain regions that were ranked in the top five for intramodular connectivity (high WMDz). In most cases, a separate set of thalamic brain regions was responsible for intermodular connectivity (high PC; see Table 2 for a full list of values for the network). Overall, the control network had more brain regions with high WMDz, high PC, or both in individual modules compared with other networks. This indicates a more interconnected network with more hub regions (Fig. 2, 3).

### The cocaine withdrawal network is driven by cortico-thalamo-hypothalamic regions

The cocaine network had 7,127 total connections and consisted of four modules, one with the majority of all brain regions and three others with a small subset of regions. In the large module (module 1; 144 brain regions), nearly one-third (32%) of the total brain regions within the module (i.e., a mixed set of midbrain-cortico-thalamic-hypothalamic-amygdalar brain regions) had high WMDz. The brain regions that drive intramodular connectivity (high WMDz) in this module did not have any intermodular connectivity (PC). Interestingly, only three brain regions in this module (subparaventricular zone, lateral posterior nucleus of the thalamus, and frontal pole cerebral cortex) reached the criterion (PC ≥ 0.30) for a high level of intermodular connectivity, suggesting sparse communication with other modules.

One of the smaller modules, a septal (triangular nucleus of the septum) and cortical (e.g., secondary motor area and dorsal anterior cingulate area) module (module 3) had a different set of thalamic brain regions that had high PC. The other two smaller modules, a prefrontal-habenular module (module 4; dorsal peduncular area [DP], induseum griseum, and lateral habenula) and a thalamic (parafascicular nucleus, mediodorsal nucleus of the thalamus, and ventral medial nucleus of the thalamus), midbrain (nucleus of the posterior commissure), and striatal (bed nucleus of the accessory olfactory tract) module (module 2) contained regions with both a high WMDz and high PC, suggesting that these regions may be potential hubs within the network. Overall, the cocaine network contained the highest number of connections in any network but had minimal interconnection between modules (Fig. 2, 4; see Table 3 for a full list of values for the network).

**Figure 4.**
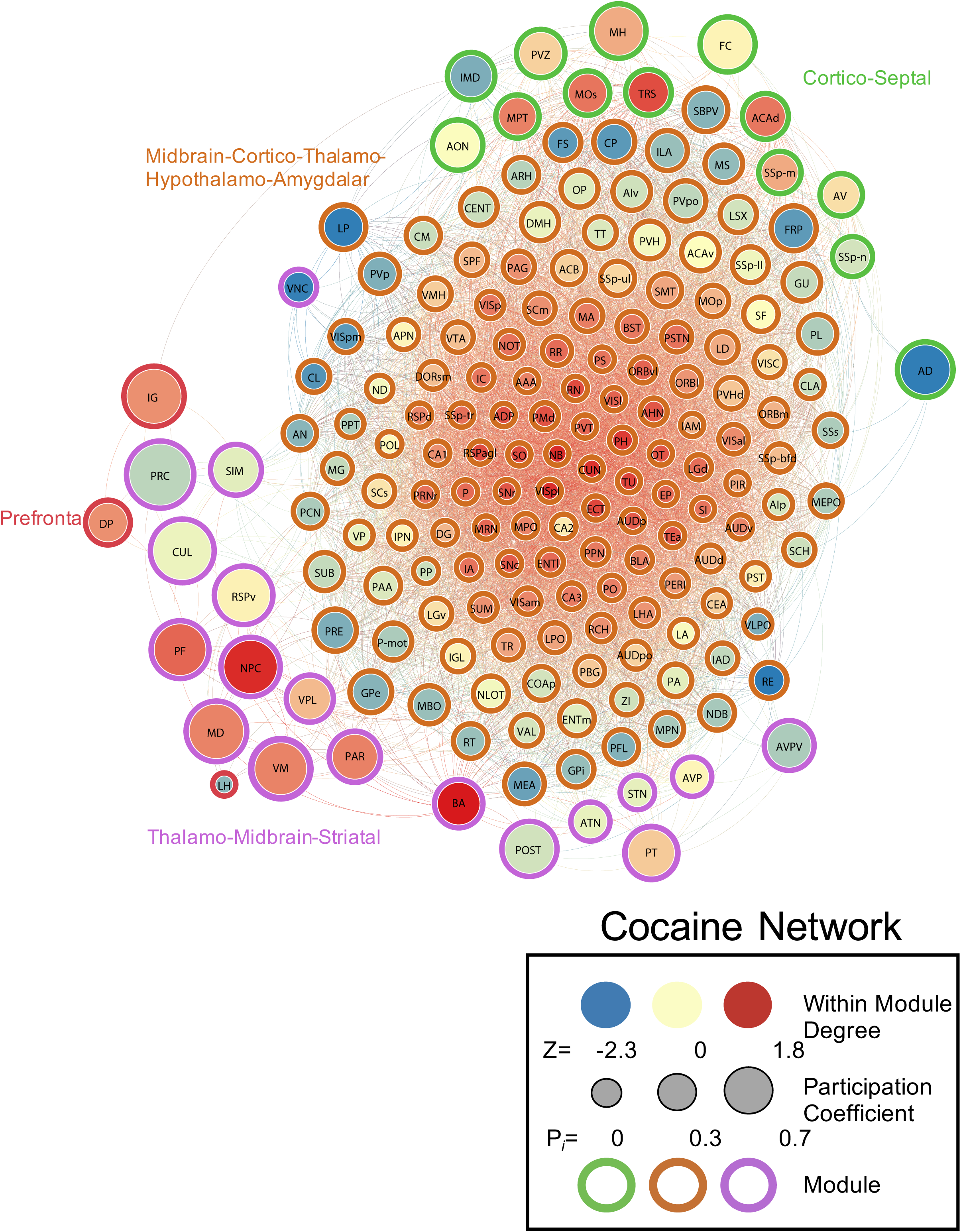
Neural network of cocaine mice during withdrawal thresholded to 0.75R. Nodes/brain regions of the network are represented by circles. The size of the node represents the participation coefficient (smaller = lower PC; larger = higher PC). The internal color of each circle represents the within-module degree Z-score (dark blue = lowest; dark red = highest). The color of the modules that are identified in Fig. 1D are represented by different colored edges. See figure key for examples of each representative component of the figure.

### The methamphetamine withdrawal network is driven by thalamic regions

The methamphetamine network had 3,182 connections and consisted of three modules, one with the majority of all brain regions and two others with a small subset of regions. In the large module (module 1), a group of thalamic (e.g., intermediodorsal nucleus of the thalamus, paraventricular nucleus of the thalamus, intergeniculate leaflet of the lateral geniculate complex, and ventral part of the lateral geniculate complex) and amygdalar (intercalated amygdala, central amygdala, and lateral amygdala) regions had high WMDz, but these brain regions did not have any intermodular connectivity (PC), and a separate set of hypothalamic, cortical, and mid/hindbrain regions was responsible for intermodular connectivity.

The second module (module 2) had several hypothalamic (e.g., mammillary body, ventrolateral preoptic nucleus, and tuberal nucleus) and pallidal (globus pallidus and internal segment) brain regions with high WMDz and a separate set of cortical regions (e.g., DP and orbital area, ventral part) and midbrain regions (e.g., posterior pretectal nucleus, nucleus of the posterior commissure, and nucleus of darkschewitsch) that had high interconnectivity with other modules (high PC).

The third module (module 3), a thalamic module, had several thalamic regions with high WMDz (e.g., ventral medial nucleus of the thalamus, posterior complex of the thalamus, parafascicular nucleus, and lateral dorsal nucleus of the thalamus). Interestingly, within this module, a separate set of thalamic regions (e.g., paracentral nucleus, ventral anterior-lateral complex of the thalamus, ventral posterior complex of the thalamus, and anterodorsal nucleus) had high PC, indicating that this module is internally directed by thalamic regions and also externally communicates through these regions. Overall, the methamphetamine network had a similar number of total connections to the control network, but it had minimal interconnections between modules (Fig. 2, 5; see Table 4 for a full list of values for the network).

**Figure 5.**
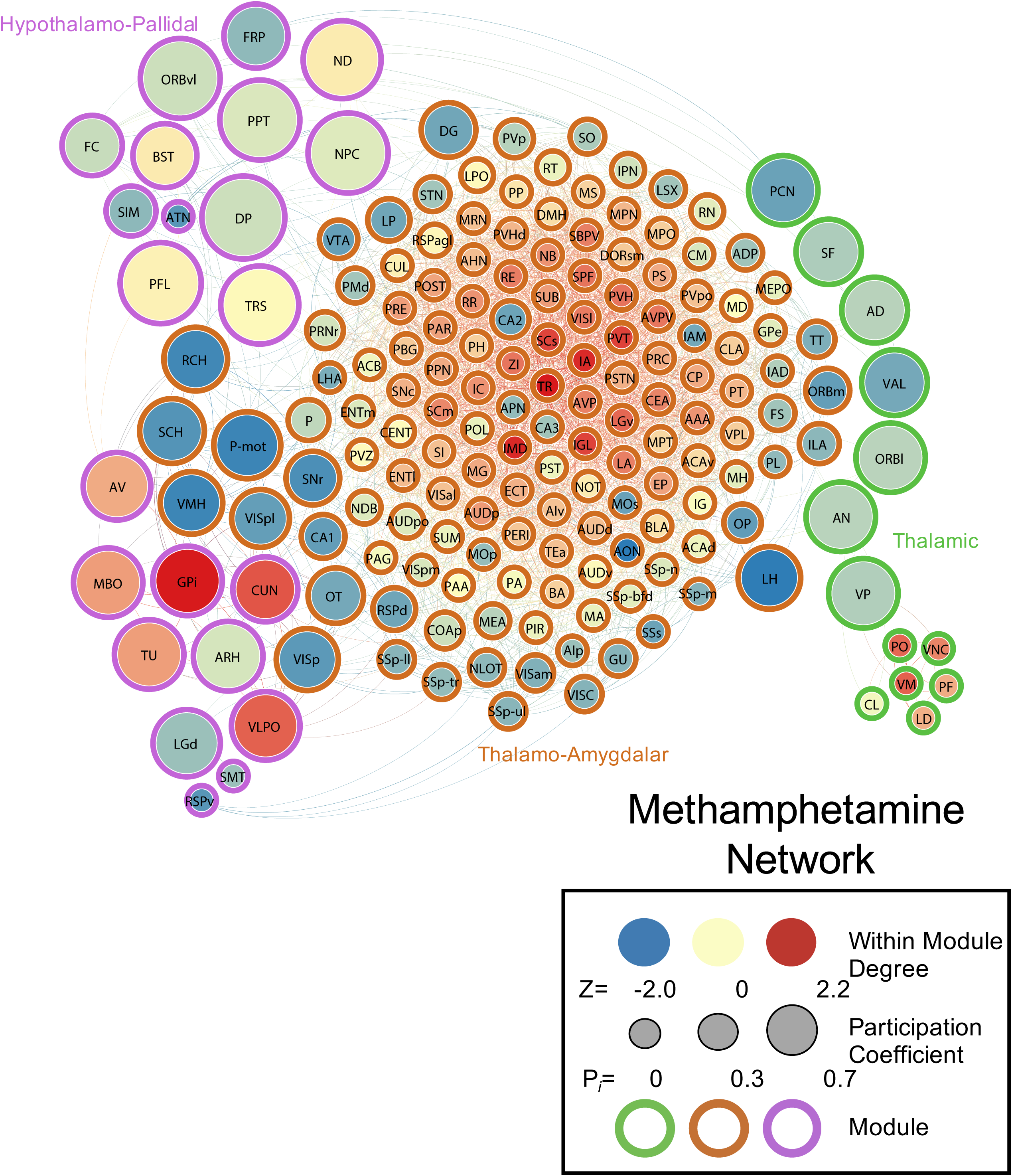
Neural network of methamphetamine mice during withdrawal thresholded to 0.75R. Nodes/brain regions of the network are represented by circles. The size of the node represents the participation coefficient (smaller = lower PC; larger = higher PC). The internal color of each circle represents the within-module degree Z-score (dark blue = lowest; dark red = highest). The color of the modules that are identified in Fig. 1E are represented by different colored edges. See figure key for examples of each representative component of the figure.

### The nicotine withdrawal network is driven by cortical and extended amygdalar regions

The nicotine network had 4,957 connections, the second most of all conditions, and consisted of five modules and one brain region (interanterodorsal nucleus of the thalamus) that was disconnected from the entire network. Overall, the nicotine network was relatively interconnected between modules and had two large modules and three medium modules.

One of the large modules (module 1) contained midbrain (e.g., pedunculopontine nucleus and periaqueductal gray), hindbrain (e.g., pons and pontine reticular nucleus), cortical (e.g., perirhinal area, posterior auditory area, ventral anterior cingulate temporal association areas, and visceral area), and subcortical (claustrum) brain regions that had high WMDz. A separate set of cortical (e.g., postsubiculum, lateral visual area, and gustatory areas), thalamic (e.g., anteroventral nucleus of the thalamus and peripeduncular nucleus), hypothalamic (e.g., posterior periventricular nucleus, supramammillary nucleus, and periventricular zone), and midbrain (e.g., midbrain reticular nucleus, ventral tegmental area, and medial pretectal area) brain regions and a few others that included the central amygdala and vestibular nuclei had high PC.

In the second large module (module 4), a set of sensory/cortical (e.g., primary somatosensory area, lower limb, ventral agranular insular area [AIv], and primary motor area) and hypothalamic (e.g., parastriatal nucleus, retrochiasmatic area, lateral preoptic area, medial preoptic area, and zona incerta) brain regions had high WMDz. All of the same sensory/cortical and hypothalamic regions had high PC and a number of other thalamic and sensory regions. Additionally, the anterior amygdalar area (AAA) also showed both high WMDz and high PC.

One of the smaller modules (module 2) consisted of hippocampal (dentate gyrus) and sensory/cortical (e.g., posterolateral visual area, anteromedial visual area, and subiculum [SUB]) regions, along with the nucleus of reuniens (RE) with high WMDz. The SUB and RE also had high PC, along with other thalamic, hypothalamic, and midbrain regions.

In another smaller module (module 3), the precommissural nucleus (PRC), medial habenula, and intergeniculate leaflet of the lateral geniculate complex (IGL) had high WMDz and high PC. Other midbrain and thalamic regions also had high PC.

In the last small module (module 5), no regions reached the criterion for high WMDz, but the orbitofrontal cortex (lateral and ventrolateral orbital area), bed nucleus of the stria terminalis, and medial amygdalar nucleus were all in the top five values (WMDz = 0.64-0.67). However, every region in this module, with the exception of the bed nucleus of the accessory olfactory tract, reached the criterion for high PC (Fig. 2, 6; see Table 5 for a full list of values for the network).

## Discussion

The present study used unbiased single-cell whole-brain imaging to identify changes in brain functional architecture after withdrawal from chronic exposure to psychostimulants. Withdrawal from psychostimulants resulted in a massive increase in neural coactivation that was associated with a decrease in modularity with varying degrees of severity, depending on the drug, compared with control mice. This decreased modularity resulted in the emergence of new network architecture and organization of the brain. Using graph theory, we identified brain regions that are most responsible for inter- and intramodular communication within each network. Withdrawal from all of the psychostimulants that were tested in the present study resulted in different network organization than the control network. The methamphetamine and cocaine withdrawal networks closely resembled each other in structural organization, primarily through thalamic motifs, whereas the nicotine withdrawal network shared some similarities with the control network. These unbiased whole-brain analyses demonstrate that psychostimulant withdrawal produces the drug-dependent remodeling of functional architecture of the brain and suggest that decreased modularity of the brain functional network may be a central feature of addiction.

We found that cocaine, methamphetamine, and nicotine withdrawal produced major increases in coordinated activity throughout the brain compared with control mice. We further found that withdrawal resulted in a decrease in modular structuring of the brain compared with control mice (seven modules). The decrease in modularity was most evident for methamphetamine withdrawal (three modules) and cocaine withdrawal (four modules), whereas nicotine withdrawal showed a smaller reduction of modularity (five modules). Such reductions of modularity are found in humans who suffer from dementia and traumatic brain injury and are associated with cognitive deficits [28–33]. Changes in network structure/functional connectivity [10–13] and cognitive function [34–36] have been observed after chronic drug use and withdrawal, suggesting that similar mechanisms may be active between these different neural disorders.

We examined the components of individual modules within each network and found that the control network was heavily driven by sensory and motor brain regions. This result confers validity to our single-cell whole-brain network analysis approach for characterizing network features because it fits with what might be expected from a normal, awake, behaving animal that explores the environment and relies heavily on sensory/motor systems. Furthermore, the control network was more interconnected between modules overall and contained several regions that could be classified as hubs of each module that are critical for network function, based on high intra- and intermodular connectivity. This suggests that the control brain may be more resilient to the disruption of function because additional hub regions may compensate more easily in response to such disruptions.

In the networks that were associated with withdrawal from psychostimulants, a shift was observed from sensory/motor regions to more subcortical (e.g., amygdalar, thalamic, hypothalamic, and midbrain) regions that drive the network. A similar effect was seen in nonhuman primates after cocaine abstinence [37], and alterations of connectivity of the somatosensory cortex are associated with smokers [38]. This may represent a shift from top-down cortical network control [39] to bottom-up subcortical network control and may reflect the greater influence of internal drives that are associated with negative affect during withdrawal in controlling the whole-brain network [40]. This shift may be a major reason why drugs are so addictive because higher cortical connectivity in humans may protect against relapse [41].

Given the modular organization of the different networks, both the control network and nicotine network had a much higher incidence of intermodular connectivity, whereas the methamphetamine and cocaine networks had only a small subset of brain regions that were connected between different modules. Similar changes in neural activity, combined with decreases in interconnectivity and network efficiency, have been observed in humans after psychostimulant use [42–44]. The nicotine network was different from the methamphetamine and cocaine networks and somewhat resembled a slightly altered control network. Similarities and differences in network properties of the three different drugs are likely to be caused by differences in receptor mechanisms and locations where each drug acts throughout the brain. Indeed, both cocaine and methamphetamine target the same dopamine transporter, whereas nicotine acts on nicotinic receptors [45–48]. These results suggest that single-cell whole-brain imaging may be used as a fingerprint or “brainprint” to characterize novel compounds by comparing whole-brain network changes to existing compounds.

The interanterodorsal nucleus of the thalamus was disconnected from the nicotine network, suggesting that it may not be involved in controlling the withdrawal network, although we cannot exclude the possibility that its disconnection may instead be a critical feature of nicotine withdrawal. One of the larger modules in the nicotine network was driven by several brain regions, two of which included the AAA and AIv, which have been suggested to be associated with nicotine withdrawal in humans [49, 50]. The methamphetamine and cocaine networks, although having distinctly different features, shared an overall motif of lower modularity and being heavily driven by thalamic brain regions. This suggests that, in a destabilized and less structured neural network, the thalamus becomes more critical to controlling the whole-brain network. The thalamus is thought to play a major role in relaying information, and the reliance of these networks on this group of regions suggests that the thalamus is not simply a relay station but has greater importance in cognitive and emotional function [51, 52]. Substantial evidence corroborates the importance of the thalamus in psychostimulant addiction and withdrawal. In a rat model of cocaine self-administration, the thalamus was found to be heavily involved in network function during acute abstinence, but changes in the network disappeared after 2 weeks [22]. Interestingly, the thalamus in humans has been shown to be hypoactive in cocaine abusers [53], and thalamic connectivity is predictive of cocaine dependence [54] and altered in infants who are exposed to cocaine [55]. Although network changes that are induced by acute withdrawal are reversed over time [22], prolonged use may lead to more permanent restructuring of the brain, and major differences between the nicotine and methamphetamine/cocaine networks may account for differences in the severity of each drug after long-term use [35, 45, 56].

In the past 40 years, the addiction field has made tremendous progress by identifying numerous brain regions that are dysregulated after psychostimulant exposure and contribute to addiction-like behaviors [3–9]. Despite this vast knowledge, however, still unclear are the ways in which these neuroadaptations, as a whole, lead to addiction. The identification of a final common brain pathway that is responsible for addiction remains elusive. The present results confirm that a substantial number of brain regions are affected by psychostimulant exposure and suggest that the final common pathway that is responsible for addiction may not reside at the level of brain regions or even single neural circuits. Instead, these results suggest that the main common phenomenon that is observed among all three of these psychostimulants is decreased modularity of whole-brain functional architecture, suggesting that the final common pathway may reside at the whole-network level. This interpretation is consistent with the literature on the modularity of complex systems, including the brain and mind, showing that lower modularity reduces the capacity of the system to adapt to its environment [57]. Such a reduction of the capacity to adapt to the environment is reminiscent of one cardinal symptom of addiction, namely continued drug use despite adverse consequences.

In summary, the present study showed that withdrawal from psychostimulants results in changes in neural network structure, including increases in coactivation among brain regions and decreases in modularity. Psychostimulant withdrawal resulted in a shift from a sensory/motor-driven network to a network that is highly driven by subcortical regions. We also found that different psychostimulants do not produce the same neural networks, although methamphetamine and cocaine shared similar properties. These findings shed light on alterations of brain function that are caused by drug exposure and identify potential brain regions that warrant future study. The present study demonstrates that psychostimulant withdrawal produces drug-dependent remodeling of the functional architecture of the brain and suggests that decreased modularity of the brain functional networks and not a specific set of brain regions may represent the final common pathway that leads to addiction. These findings may prove critical to designing future treatment approaches for drug abuse.

## Materials and Methods

### Animals

Male C57BL/6J mice were bred at The Scripps Research Institute. They were 20-30 g and 60 days old at the start of the experiment. The mice were maintained on a 12 h/12 h light/dark cycle with *ad libitum* access to food and water. All of the procedures were conducted in strict adherence to the National Institutes of Health Guide for the Care and Use of Laboratory Animals and approved by The Scripps Research Institute Institutional Animal Care and Use Committee.

### Drugs

The doses were 4 mg/kg/day for methamphetamine, 24 mg/kg/day for nicotine, and 60 mg/kg/day for cocaine. These doses were chosen based on previous studies that indicated rewarding effects during use, resulting in withdrawal-like symptoms after the cessation of chronic use [14–20]. Each drug was dissolved in saline, and the pH was adjusted to 7.4. The drugs were loaded into osmotic minipumps (Alzet; model no. 1002). The minipumps sat overnight in saline before insertion to ensure that drug delivery would begin immediately.

### Minipump implantation and removal

The mice were split into four groups for the experiment: methamphetamine withdrawal group (*n* = 5), nicotine withdrawal group (*n* = 5), cocaine withdrawal group (*n* = 5), and saline control group (*n* = 4). Each mouse was surgically implanted with an osmotic minipump for methamphetamine, nicotine, cocaine, and saline based on group assignment. The minipumps were implanted in the lower back of each mouse under anesthesia. After brief recovery, the mice were returned to their home cages. The mice remained in their home cages for 1 week to allow for chronic infusion of the drug.

After 1 week, the minipumps were surgically removed under anesthesia to allow for drug washout and withdrawal to begin. Mice in the nicotine, cocaine, and saline groups were perfused 8 h after removal of the minipumps. Mice in the methamphetamine group were perfused 12 h after removal of the minipumps. These time points were chosen to represent an acute withdrawal period from each drug (e.g., a minimum of 4 h without the drug present) and based on the half-life of each drug in mice [58–62].

### Tissue collection

The mice were deeply anesthetized and perfused with 15 ml of phosphate-buffered saline (PBS) followed by 50 ml of 4% formaldehyde. The brains were postfixed in formaldehyde overnight. The next day, the brains were washed for 30 min three times with PBS and transferred to a PBS/0.1% azide solution at 4°C for 2-3 days before processing via iDISCO+.

### iDISCO+

The iDISCO+ procedure was performed as reported by Renier et al. [63, 64].

### Immunostaining

Fixed samples were washed in 20% methanol (in double-distilled H_2_O) for 1 h, 40% methanol for 1 h, 60% methanol for 1 h, 80% methanol for 1 h, and 100% methanol for 1 h twice. The samples were then precleared with overnight incubation in 33% methanol/66% dichloromethane (DCM; Sigma, catalog no. 270997-12X100ML). The next day, the samples were bleached with 5% H_2_O_2_ (1 volume of 30% H_2_O_2_ for 5 volumes of methanol, ice cold) at 4°C overnight. After bleaching, the samples were slowly re-equilibrated at room temperature and rehydrated in 80% methanol in double-distilled H_2_O for 1 h, 60% methanol for 1 h, 40% methanol for 1 h, 20% methanol for 1 h, PBS for 1 h, and PBS/0.2% TritonX-100 for 1 h twice. The samples were then incubated in PBS/0.2% TritonX-100/20% dimethylsulfoxide (DMSO)/0.3 M glycine at 37°C for 2 days and then blocked in PBS/0.2% TritonX-100/10% DMSO/6% donkey serum at 37°C for 2 days. The samples were then incubated in rabbit anti c-*fos* (1:2000; Synaptic Systems catalog number 226 003) in PBS-0.2% Tween with 10 μg/ml heparin (PTwH)/5% DMSO/3% donkey serum at 37°C for 7 days. The samples were then washed in PTwH for 24 h (five changes of the PTwH solution over that time) and incubated in donkey anti-rabbit Alexa647 (1:500; Invitrogen, catalog no. A31573) in PTwH/3% donkey serum at 37°C for 7 days. The samples were finally washed in PTwH for 1 day before clearing and imaging.

### Sample clearing

Immunolabeled brains were cleared using the procedure of Reiner et al. (2016) [63]. The samples were dehydrated in 20% methanol in double-distilled H_2_O for 1 h, 40% methanol for 1 h, 60% methanol for 1 h, 80% methanol for 1 h, 100% methanol for 1 h, and 100% methanol again overnight. The next day, the samples were incubated for 3 h in 33% methanol/66% DCM until they sank to the bottom of the incubation tube. The methanol was then washed for 20 min twice in 100% DCM. Finally, the samples were incubated in dibenzyl ether (DBE; Sigma, catalog no. 108014-1KG) until clear and then stored in DBE at room temperature until imaged.

### Image acquisition

Left hemispheres of cleared samples were imaged in the sagittal orientation (right lateral side up) on a light-sheet microscope (Ultramicroscope II, LaVision Biotec) equipped with an sCMOS camera (Andor Neo) and 2×/0.5 objective lens (MVPLAPO 2×) equipped with a 6 mm working distance dipping cap. Imspector Microscope controller v144 software was used. The microscope was equipped with an NKT Photonics SuperK EXTREME EXW-12 white light laser with three fixed light sheet generating lenses on each side. Scans were made at 0.8× magnification (1.6× effective magnification) with a light sheet numerical aperture of 0.148. Excitation filters of 480/30, 560/40, and 630/30 nm were used. Emission filters of 525/50, 595/40, and 680/30 nm were used. The samples were scanned with a step size of 3 μm using dynamic horizontal scanning from one side (the right) for the 560 and 630 nm channels (20 acquisitions per plane with 240 ms exposure, combined into one image using the horizontal adaptive algorithm) and without horizontal scanning for the 480 nm channel using two-sided illumination (100 ms exposure for each side, combined into one image using the blending algorithm). To accelerate acquisition, both channels where acquired in two separate scans. To account for micro-movements of the samples that may occur between scans, three-dimensional image affine registration was performed to align both channels using ClearMap [63].

### Data analysis

#### Identification of activated brain regions

Images that were acquired from the light-sheet microscope were analyzed from the end of the olfactory bulbs (the olfactory bulbs were not included in the analysis) to the beginning of the hindbrain and cerebellum. Counts of Fos-positive nuclei from each sample were identified for each brain region using ClearMap [63]. ClearMap uses autofluorescence that is acquired in the 488 nm channel to align the brain to the Allen Mouse Brain Atlas [65] and then registers Fos counts to regions that are annotated by the atlas. The data were normalized to a log_10_ value to reduce variability and bring brain regions with high numbers (e.g., thousands) and low numbers (e.g., tens to hundreds) of Fos counts to a similar scale.

#### Identification of co-activation within individual networks

Separate inter-regional Pearson correlations were then calculated using Statistica software (Tibco) across animals in the saline, cocaine, methamphetamine, and nicotine groups to compare the log_10_ Fos data from each brain region to each of the other brain regions. See Table 1 for a list of brain regions, their abbreviations, and their Allen atlas grouping.

### Hierarchical clustering

Previous rat and mouse studies that examined functional connectivity used 5-8 animals [21, 22]. The number of samples that are examined in functional connectivity studies is the number of potential connections (i.e., 178 total brain regions all connecting with each other for each treatment). Furthermore, hierarchical clustering organizes brain regions into modules by grouping regions that show a similar coactivation profile across all other brain regions. Thus, more total connections minimize the effect that an inaccurate brain region-to-brain region connection has on network organization and overall network structure.

Inter-regional Fos correlations were then used to calculate complete Euclidean distances between each pair of brain regions in each group of mice. The distance matrices were then hierarchically clustered using R Studio software by both row and column using the complete method to identify modules of coactivation within each treatment group. The hierarchical cluster dendrograms were trimmed at half the height of each given tree to split the dendrogram into specific modules. The result of a decrease in modularity that is attributable to psychostimulant use was consistent across multiple tree-cutting thresholds (Fig. 1B).

### Graph theory identification of functional networks

We used a graph theory-based approach to identify the functional neural networks that were associated with each treatment condition. Graph theory is a branch of mathematics that is used to analyze complex networks, such as social, financial, protein, and neural networks [21, 66–77]. Using graph theory, functional networks can be delineated, and key brain regions of the network can be identified [21, 69, 78, 79].

Previous studies of regional connectivity profiles in Fos coactivation networks have focused on global measures of connectivity (e.g., degree) [21]. However, in correlation-based networks, these measures can be strongly influenced by the size of the subnetwork (module) in which a node participates [80]. For the graph theory analyses, we were interested in regional properties and not module size *per se*. Thus, module structure needs to be considered when examining the role that each region plays in the network. To accomplish this, we utilized two widely used centrality metrics that were designed for application to modular systems. The WMDz indexes the relative importance of a region within its own module (e.g., intramodule connectivity), and the PC indexes the extent to which a region connects diversely to multiple modules (e.g., intermodule connectivity) [27].

We used the Pearson correlation values that were calculated for the brain regions from each treatment. Prior to plotting and calculating regional connectivity metrics, the network was thresholded to remove any edges that were weaker than *R* = 0.75. As such, visualization and graph theory analyses were performed using only edges with positive weights. Regional connectivity metrics (PC and WMDz) were calculated as originally defined by Guimerà and Amaral (2005) [27], modified for application to networks with weighted edges. PC and WMDz were calculated using a customized version of the bctpy Python package (https://github.com/aestrivex/bctpy), which is derived from the MATLAB implementation of Brain Connectivity Toolbox [78].

For WMDz, let *k_i_* (within-module degree) be the summed weight of all edges between region *i* and other regions in module *s*_*i*_. Then, 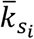 is the average within-module degree of all regions in module *s*_*i*_, and 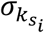 is the standard deviation of those values. The Z-scored version of within-module degree (WMDz) is then defined as:

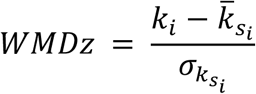

This provides a measure of the extent to which each region is connected to other regions in the

For PC, let *k_is_* (between-module degree) be the summed weight of all edges between region *i* and regions in module *s*, and let *k_i_* (total degree) be the summed weight of all edges between region *i* and all other regions in the network. The PC of each region is then defined as:

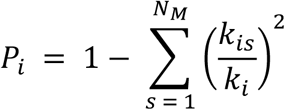

This provides a measure of the extent to which the connections of a region are distributed mostly within its own module (PC approaching 0) or distributed evenly among all modules (PC approaching 1).

A high PC was considered ≥ 0.30, and a high WMDz was considered ≥ 0.80. Previous studies have used ranges of ≥ 0.30-0.80 for high PC and ≥ 1.5-2.5 for high WMDz [27, 77]. Because of differences in the sizes/types of networks that were examined and the methods that were used (e.g., Fos *vs*. functional magnetic resonance imaging), we adjusted the range for consideration as having high PC and WMDz accordingly.

Network visualization was performed using a combination of Gephi 0.9.2 software [81] and Adobe Illustrator software. Nodes were positioned using the Force Atlas 2 algorithm [82] with a handful of nodes that were repositioned manually for better visual organization.

