## Supplementary figures and images for "Characterization of the brain functional architecture of psychostimulant withdrawal using single-cell whole brain imaging"

### Supplemental Figure 1

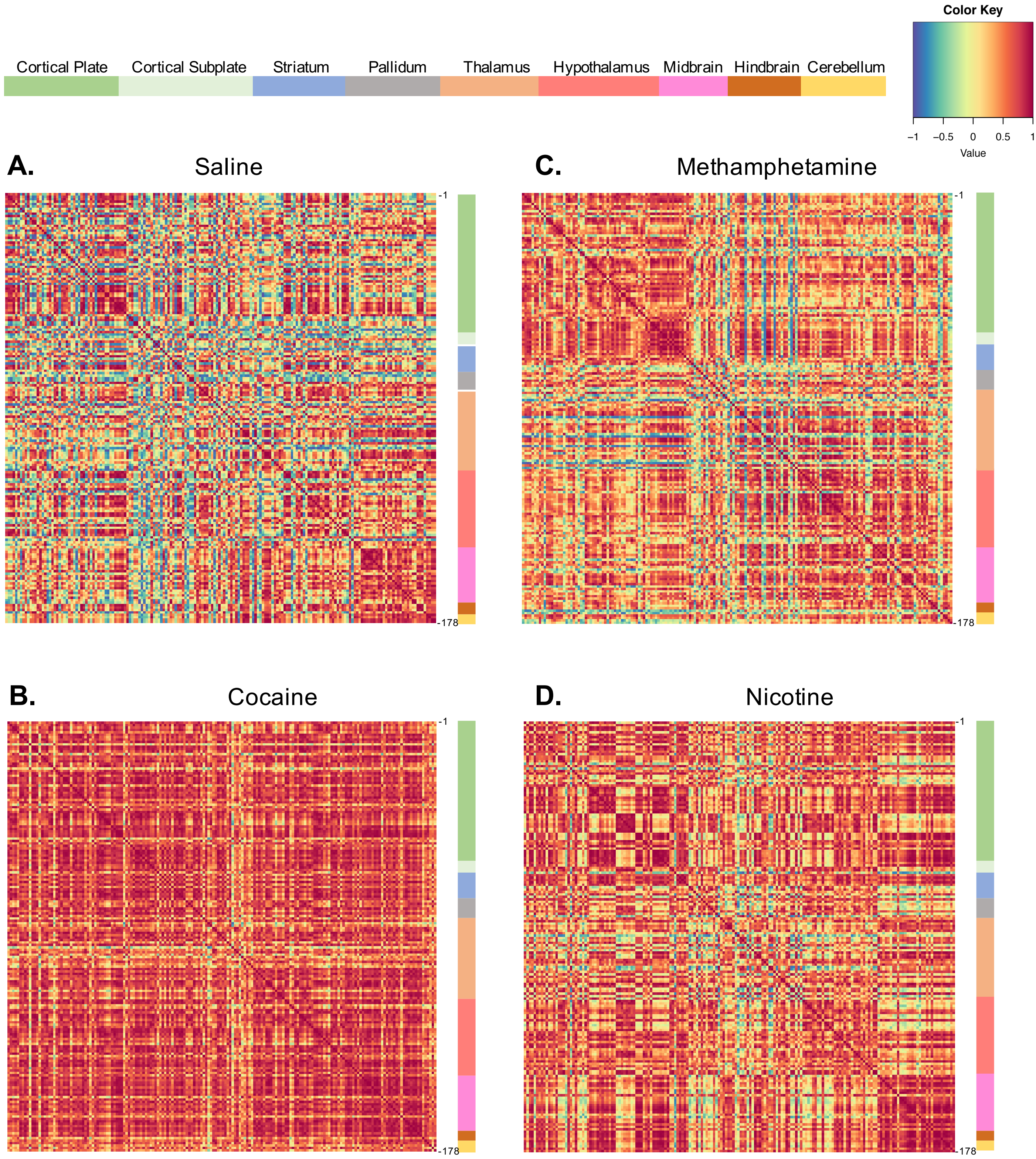
