## Supplemental Table 1 for "Characterization of the brain functional architecture of psychostimulant withdrawal using single-cell whole brain imaging"

**Supplementary Table 1: Brain regions list**

| **Brain Region** | **Abbreviation** | **Allen Group Name** |
| --- | --- | --- |
| Agranular insular area posterior part | AIp | Cortical Plate |
| Agranular insular area ventral part | AIv | Cortical Plate |
| Anterior cingulate area dorsal part | ACAd | Cortical Plate |
| Anterior cingulate area ventral part | ACAv | Cortical Plate |
| Anterior olfactory nucleus | AON | Cortical Plate |
| Anterolateral visual area | VISal | Cortical Plate |
| Anteromedial visual area | VISam | Cortical Plate |
| Cortical amygdalar area posterior part | COAp | Cortical Plate |
| Dentate gyrus | DG | Cortical Plate |
| Dorsal auditory area | AUDd | Cortical Plate |
| Dorsal peduncular area | DP | Cortical Plate |
| Ectorhinal area | ECT | Cortical Plate |
| Entorhinal area lateral part | ENTl | Cortical Plate |
| Entorhinal area medial part | ENTm | Cortical Plate |
| Fasciola cinerea | FC | Cortical Plate |
| Field CA1 | CA1 | Cortical Plate |
| Field CA2 | CA2 | Cortical Plate |
| Field CA3 | CA3 | Cortical Plate |
| Frontal pole cerebral cortex | FRP | Cortical Plate |
| Gustatory areas | GU | Cortical Plate |
| Induseum griseum | IG | Cortical Plate |
| Infralimbic area | ILA | Cortical Plate |
| Lateral visual area | VISl | Cortical Plate |
| Nucleus of the lateral olfactory tract | NLOT | Cortical Plate |
| Orbital area lateral part | ORBl | Cortical Plate |
| Orbital area medial part | ORBm | Cortical Plate |
| Orbital area ventrolateral part | ORBvl | Cortical Plate |
| Parasubiculum | PAR | Cortical Plate |
| Perirhinal area | PERI | Cortical Plate |
| Piriform area | PIR | Cortical Plate |
| Piriform-amygdalar area | PAA | Cortical Plate |
| Posterior auditory area | AUDpo | Cortical Plate |
| Posterolateral visual area | VISpl | Cortical Plate |
| Posteromedial visual area | VISpm | Cortical Plate |
| Postpiriform transition area | TR | Cortical Plate |
| Postsubiculum | POST | Cortical Plate |
| Prelimbic area | PL | Cortical Plate |
| Presubiculum | PRE | Cortical Plate |
| Primary auditory area | AUDp | Cortical Plate |
| Primary motor area | MOp | Cortical Plate |
| Primary somatosensory area barrel field | SSp-bfd | Cortical Plate |
| Primary somatosensory area lower limb | SSp-ll | Cortical Plate |
| Primary somatosensory area mouth | SSp-m | Cortical Plate |
| Primary somatosensory area nose | SSp-n | Cortical Plate |
| Primary somatosensory area trunk | SSp-tr | Cortical Plate |
| Primary somatosensory area upper limb | SSp-ul | Cortical Plate |
| Primary visual area | VISp | Cortical Plate |
| Retrosplenial area dorsal part | RSPd | Cortical Plate |
| Retrosplenial area lateral agranular part | RSPagl | Cortical Plate |
| Retrosplenial area ventral part | RSPv | Cortical Plate |
| Secondary motor area | MOs | Cortical Plate |
| Subiculum | SUB | Cortical Plate |
| Supplemental somatosensory area | SSs | Cortical Plate |
| Taenia tecta | TT | Cortical Plate |
| Temporal association areas | TEa | Cortical Plate |
| Ventral auditory area | AUDv | Cortical Plate |
| Visceral area | VISC | Cortical Plate |
| Basolateral amygdalar nucleus | BLA | Cortical Subplate |
| Claustrum | CLA | Cortical Subplate |
| Endopiriform nucleus | EP | Cortical Subplate |
| Lateral amygdalar nucleus | LA | Cortical Subplate |
| Posterior amygdalar nucleus | PA | Cortical Subplate |
| Anterior amygdalar area | AAA | Striatum |
| Bed nucleus of the accessory olfactory tract | BA | Striatum |
| Caudoputamen | CP | Striatum |
| Central amygdalar nucleus | CEA | Striatum |
| Fundus of striatum | FS | Striatum |
| Intercalated amygdalar nucleus | IA | Striatum |
| Lateral septal complex | LSX | Striatum |
| Medial amygdalar nucleus | MEA | Striatum |
| Nucleus accumbens | ACB | Striatum |
| Olfactory tubercle | OT | Striatum |
| Septofimbrial nucleus | SF | Striatum |
| Bed nuclei of the stria terminalis | BST | Pallidum |
| Diagonal band nucleus | NDB | Pallidum |
| Globus pallidus external segment | GPe | Pallidum |
| Globus pallidus internal segment | GPi | Pallidum |
| Magnocellular nucleus | MA | Pallidum |
| Medial septal nucleus | MS | Pallidum |
| Substantia innominata | SI | Pallidum |
| Triangular nucleus of septum | TRS | Pallidum |
| Anterior group of the dorsal thalamus | ATN | Thalamus |
| Anterodorsal nucleus | AD | Thalamus |
| Anteroventral nucleus of thalamus | AV | Thalamus |
| Central lateral nucleus of the thalamus | CL | Thalamus |
| Central medial nucleus of the thalamus | CM | Thalamus |
| Dorsal part of the lateral geniculate complex | LGd | Thalamus |
| Interanterodorsal nucleus of the thalamus | IAD | Thalamus |
| Interanteromedial nucleus of the thalamus | IAM | Thalamus |
| Intergeniculate leaflet of the lateral geniculate complex | IGL | Thalamus |
| Intermediodorsal nucleus of the thalamus | IMD | Thalamus |
| Lateral dorsal nucleus of thalamus | LD | Thalamus |
| Lateral habenula | LH | Thalamus |
| Lateral posterior nucleus of the thalamus | LP | Thalamus |
| Medial geniculate complex | MG | Thalamus |
| Medial habenula | MH | Thalamus |
| Mediodorsal nucleus of thalamus | MD | Thalamus |
| Nucleus of reuniens | RE | Thalamus |
| Paracentral nucleus | PCN | Thalamus |
| Parafascicular nucleus | PF | Thalamus |
| Parataenial nucleus | PT | Thalamus |
| Paraventricular nucleus of the thalamus | PVT | Thalamus |
| Peripeduncular nucleus | PP | Thalamus |
| Posterior complex of the thalamus | PO | Thalamus |
| Posterior limiting nucleus of the thalamus | POL | Thalamus |
| Reticular nucleus of the thalamus | RT | Thalamus |
| Submedial nucleus of the thalamus | SMT | Thalamus |
| Subparafascicular nucleus | SPF | Thalamus |
| Thalamus sensory-motor cortex related | DORsm | Thalamus |
| Ventral anterior-lateral complex of the thalamus | VAL | Thalamus |
| Ventral medial nucleus of the thalamus | VM | Thalamus |
| Ventral part of the lateral geniculate complex | LGv | Thalamus |
| Ventral posterior complex of the thalamus | VP | Thalamus |
| Ventral posterolateral nucleus of the thalamus | VPL | Thalamus |
| Anterior hypothalamic nucleus | AHN | Hypothalamus |
| Anterodorsal preoptic nucleus | ADP | Hypothalamus |
| Anteroventral periventricular nucleus | AVPV | Hypothalamus |
| Anteroventral preoptic nucleus | AVP | Hypothalamus |
| Arcuate hypothalamic nucleus | ARH | Hypothalamus |
| Dorsal premammillary nucleus | PMd | Hypothalamus |
| Dorsomedial nucleus of the hypothalamus | DMH | Hypothalamus |
| Lateral hypothalamic area | LHA | Hypothalamus |
| Lateral preoptic area | LPO | Hypothalamus |
| Mammillary body | MBO | Hypothalamus |
| Medial preoptic area | MPO | Hypothalamus |
| Medial preoptic nucleus | MPN | Hypothalamus |
| Median preoptic nucleus | MEPO | Hypothalamus |
| Parastrial nucleus | PS | Hypothalamus |
| Parasubthalamic nucleus | PSTN | Hypothalamus |
| Paraventricular hypothalamic nucleus | PVH | Hypothalamus |
| Paraventricular hypothalamic nucleus descending division | PVHd | Hypothalamus |
| Periventricular hypothalamic nucleus posterior part | PVp | Hypothalamus |
| Periventricular hypothalamic nucleus preoptic part | PVpo | Hypothalamus |
| Periventricular zone | PVZ | Hypothalamus |
| Posterior hypothalamic nucleus | PH | Hypothalamus |
| Preparasubthalamic nucleus | PST | Hypothalamus |
| Retrochiasmatic area | RCH | Hypothalamus |
| Subparaventricular zone | SBPV | Hypothalamus |
| Subthalamic nucleus | STN | Hypothalamus |
| Suprachiasmatic nucleus | SCH | Hypothalamus |
| Supramammillary nucleus | SUM | Hypothalamus |
| Supraoptic nucleus | SO | Hypothalamus |
| Tuberal nucleus | TU | Hypothalamus |
| Ventrolateral preoptic nucleus | VLPO | Hypothalamus |
| Ventromedial hypothalamic nucleus | VMH | Hypothalamus |
| Zona incerta | ZI | Hypothalamus |
| Anterior pretectal nucleus | APN | Midbrain |
| Cuneiform nucleus | CUN | Midbrain |
| Inferior colliculus | IC | Midbrain |
| Interpeduncular nucleus | IPN | Midbrain |
| Medial pretectal area | MPT | Midbrain |
| Midbrain reticular nucleus | MRN | Midbrain |
| Midbrain reticular nucleus retrorubral area | RR | Midbrain |
| Nucleus of Darkschewitsch | ND | Midbrain |
| Nucleus of the brachium of the inferior colliculus | NB | Midbrain |
| Nucleus of the optic tract | NOT | Midbrain |
| Nucleus of the posterior commissure | NPC | Midbrain |
| Olivary pretectal nucleus | OP | Midbrain |
| Parabigeminal nucleus | PBG | Midbrain |
| Pedunculopontine nucleus | PPN | Midbrain |
| Periaqueductal gray | PAG | Midbrain |
| Posterior pretectal nucleus | PPT | Midbrain |
| Precommissural nucleus | PRC | Midbrain |
| Red nucleus | RN | Midbrain |
| Substantia nigra compact part | SNc | Midbrain |
| Substantia nigra reticular part | SNr | Midbrain |
| Superior colliculus motor related | SCm | Midbrain |
| Superior colliculus sensory related | SCs | Midbrain |
| Ventral tegmental area | VTA | Midbrain |
| Pons | P | Hindbrain |
| Pons motor related | P-mot | Hindbrain |
| Pontine reticular nucleus | PRNr | Hindbrain |
| Vestibular nuclei | VNC | Hindbrain |
| Ansiform lobule | AN | Cerebellum |
| Central lobule | CENT | Cerebellum |
| Culmen | CUL | Cerebellum |
| Paraflocculus | PFL | Cerebellum |
| Simple lobule | SIM | Cerebellum |
