## Supplemental Table 5 for "Characterization of the brain functional architecture of psychostimulant withdrawal using single-cell whole brain imaging"

**Supplementary Table 5: Nicotine Network Values**

| Region | Module | PC | WMDz |
| --- | --- | --- | --- |
| Agranular insular area posterior part | 1 | 0.03 | 0.12 |
| Agranular insular area ventral part | 4 | 0.43 | 0.85 |
| Anterior cingulate area dorsal part | 1 | 0.17 | 0.87 |
| Anterior cingulate area ventral part | 1 | 0.22 | 0.94 |
| Anterior olfactory nucleus | 5 | 0.42 | 0.27 |
| Anterolateral visual area | 1 | 0.38 | -0.12 |
| Anteromedial visual area | 2 | 0.57 | 0.97 |
| Cortical amygdalar area posterior part | 1 | 0.12 | -0.46 |
| Dentate gyrus | 2 | 0.58 | 0.99 |
| Dorsal auditory area | 1 | 0.20 | 0.52 |
| Dorsal peduncular area | 4 | 0.49 | 0.33 |
| Ectorhinal area | 1 | 0.13 | 0.78 |
| Entorhinal area lateral part | 1 | 0.28 | -1.50 |
| Entorhinal area medial part | 4 | 0.57 | -0.12 |
| Fasciola cinerea | 1 | 0.25 | -0.35 |
| Field CA1 | 4 | 0.34 | -0.11 |
| Field CA2 | 4 | 0.36 | 0.33 |
| Field CA3 | 4 | 0.44 | 0.80 |
| Frontal pole cerebral cortex | 5 | 0.49 | -2.12 |
| Gustatory areas | 1 | 0.44 | -0.40 |
| Induseum griseum | 3 | 0.47 | -1.01 |
| Infralimbic area | 4 | 0.42 | 0.66 |
| Lateral visual area | 1 | 0.38 | 0.27 |
| Nucleus of the lateral olfactory tract | 4 | 0.41 | -1.16 |
| Orbital area lateral part | 5 | 0.44 | 0.67 |
| Orbital area medial part | 4 | 0.48 | -2.31 |
| Orbital area ventrolateral part | 5 | 0.46 | 0.66 |
| Parasubiculum | 4 | 0.59 | 0.18 |
| Perirhinal area | 1 | 0.13 | 1.02 |
| Piriform area | 1 | 0.00 | 0.11 |
| Piriform-amygdalar area | 1 | 0.11 | -2.62 |
| Posterior auditory area | 1 | 0.12 | 0.95 |
| Posterolateral visual area | 2 | 0.55 | 0.98 |
| Posteromedial visual area | 1 | 0.38 | -0.12 |
| Postpiriform transition area | 1 | 0.10 | 0.58 |
| Postsubiculum | 1 | 0.34 | 0.58 |
| Prelimbic area | 1 | 0.16 | -1.81 |
| Presubiculum | 1 | 0.27 | 0.69 |
| Primary auditory area | 1 | 0.07 | 0.85 |
| Primary motor area | 4 | 0.43 | 0.82 |
| Primary somatosensory area barrel field | 5 | 0.43 | 0.58 |
| Primary somatosensory area lower limb | 4 | 0.44 | 1.12 |
| Primary somatosensory area mouth | 4 | 0.61 | -0.75 |
| Primary somatosensory area nose | 4 | 0.54 | 0.67 |
| Primary somatosensory area trunk | 4 | 0.48 | 0.70 |
| Primary somatosensory area upper limb | 4 | 0.44 | 0.26 |
| Primary visual area | 4 | 0.50 | 0.56 |
| Retrosplenial area dorsal part | 1 | 0.26 | 0.30 |
| Retrosplenial area lateral agranular part | 1 | 0.21 | 0.67 |
| Retrosplenial area ventral part | 1 | 0.20 | 0.41 |
| Secondary motor area | 4 | 0.55 | 0.27 |
| Subiculum | 2 | 0.66 | 0.97 |
| Supplemental somatosensory area | 1 | 0.22 | 0.73 |
| Taenia tecta | 4 | 0.53 | 0.65 |
| Temporal association areas | 1 | 0.14 | 0.93 |
| Ventral auditory area | 1 | 0.09 | 0.85 |
| Visceral area | 1 | 0.09 | 0.90 |
| Basolateral amygdalar nucleus | 1 | 0.25 | 0.71 |
| Claustrum | 1 | 0.18 | 1.06 |
| Endopiriform nucleus | 1 | 0.10 | -0.96 |
| Lateral amygdalar nucleus | 1 | 0.24 | 0.36 |
| Posterior amygdalar nucleus | 4 | 0.35 | 0.54 |
| Anterior amygdalar area | 4 | 0.52 | 0.86 |
| Bed nucleus of the accessory olfactory tract | 5 | 0.13 | -0.68 |
| Caudoputamen | 2 | 0.61 | -0.17 |
| Central amygdalar nucleus | 1 | 0.36 | -0.01 |
| Fundus of striatum | 2 | 0.59 | 0.47 |
| Intercalated amygdalar nucleus | 1 | 0.29 | 0.46 |
| Lateral septal complex | 1 | 0.22 | -1.88 |
| Medial amygdalar nucleus | 5 | 0.49 | 0.64 |
| Nucleus accumbens | 4 | 0.39 | 0.70 |
| Olfactory tubercle | 4 | 0.39 | -2.16 |
| Septofimbrial nucleus | 1 | 0.16 | -0.31 |
| Bed nuclei of the stria terminalis | 5 | 0.49 | 0.65 |
| Diagonal band nucleus | 5 | 0.44 | -0.33 |
| Globus pallidus external segment | 4 | 0.49 | -2.31 |
| Globus pallidus internal segment | 4 | 0.45 | 0.27 |
| Magnocellular nucleus | 4 | 0.41 | 0.43 |
| Medial septal nucleus | 2 | 0.66 | 0.48 |
| Substantia innominata | 5 | 0.46 | 0.14 |
| Triangular nucleus of septum | 3 | 0.49 | 0.28 |
| Anterior group of the dorsal thalamus | 4 | 0.60 | -1.42 |
| Anterodorsal nucleus | 3 | 0.53 | -1.16 |
| Anteroventral nucleus of thalamus | 1 | 0.53 | -2.45 |
| Central lateral nucleus of the thalamus | 2 | 0.67 | -1.12 |
| Central medial nucleus of the thalamus | 1 | 0.17 | 0.09 |
| Dorsal part of the lateral geniculate complex | 4 | 0.00 | -2.07 |
| Interanteromedial nucleus of the thalamus | 1 | 0.27 | -2.07 |
| Intergeniculate leaflet of the lateral geniculate complex | 3 | 0.50 | 1.16 |
| Intermediodorsal nucleus of the thalamus | 1 | 0.12 | -0.71 |
| Lateral dorsal nucleus of thalamus | 5 | 0.47 | 0.64 |
| Lateral habenula | 3 | 0.47 | 0.75 |
| Lateral posterior nucleus of the thalamus | 5 | 0.44 | -2.94 |
| Medial geniculate complex | 1 | 0.22 | 0.81 |
| Medial habenula | 3 | 0.38 | 1.19 |
| Mediodorsal nucleus of thalamus | 1 | 0.23 | 0.03 |
| Nucleus of reuniens | 2 | 0.67 | 0.86 |
| Paracentral nucleus | 4 | 0.58 | 0.66 |
| Parafascicular nucleus | 2 | 0.68 | -0.14 |
| Parataenial nucleus | 3 | 0.40 | -0.61 |
| Paraventricular nucleus of the thalamus | 3 | 0.46 | -1.06 |
| Peripeduncular nucleus | 1 | 0.41 | -1.81 |
| Posterior complex of the thalamus | 5 | 0.45 | 0.26 |
| Posterior limiting nucleus of the thalamus | 1 | 0.21 | -0.13 |
| Reticular nucleus of the thalamus | 4 | 0.25 | -0.76 |
| Submedial nucleus of the thalamus | 4 | 0.45 | 0.72 |
| Subparafascicular nucleus | 1 | 0.25 | 0.99 |
| Thalamus sensory-motor cortex related | 1 | 0.20 | -0.83 |
| Ventral anterior-lateral complex of the thalamus | 5 | 0.46 | 0.27 |
| Ventral medial nucleus of the thalamus | 4 | 0.45 | 0.06 |
| Ventral part of the lateral geniculate complex | 1 | 0.21 | 0.44 |
| Ventral posterior complex of the thalamus | 5 | 0.48 | 0.26 |
| Ventral posterolateral nucleus of the thalamus | 1 | 0.12 | -2.84 |
| Anterior hypothalamic nucleus | 5 | 0.44 | 0.51 |
| Anterodorsal preoptic nucleus | 4 | 0.40 | 0.58 |
| Anteroventral periventricular nucleus | 4 | 0.38 | -0.22 |
| Anteroventral preoptic nucleus | 4 | 0.30 | 0.05 |
| Arcuate hypothalamic nucleus | 2 | 0.49 | -0.42 |
| Dorsal premammillary nucleus | 2 | 0.64 | 0.50 |
| Dorsomedial nucleus of the hypothalamus | 2 | 0.61 | 0.75 |
| Lateral hypothalamic area | 4 | 0.44 | -0.06 |
| Lateral preoptic area | 4 | 0.40 | 0.85 |
| Mammillary body | 1 | 0.39 | -0.30 |
| Medial preoptic area | 4 | 0.40 | 0.81 |
| Medial preoptic nucleus | 5 | 0.45 | 0.50 |
| Median preoptic nucleus | 4 | 0.29 | -1.51 |
| Parastrial nucleus | 4 | 0.45 | 1.11 |
| Parasubthalamic nucleus | 2 | 0.42 | -0.99 |
| Paraventricular hypothalamic nucleus | 1 | 0.22 | 0.88 |
| Paraventricular hypothalamic nucleus descending division | 2 | 0.65 | 0.72 |
| Periventricular hypothalamic nucleus posterior part | 1 | 0.34 | 0.21 |
| Periventricular hypothalamic nucleus preoptic part | 4 | 0.21 | -0.74 |
| Periventricular zone | 1 | 0.35 | -0.50 |
| Posterior hypothalamic nucleus | 1 | 0.28 | 0.76 |
| Preparasubthalamic nucleus | 4 | 0.45 | -2.20 |
| Retrochiasmatic area | 4 | 0.45 | 0.98 |
| Subparaventricular zone | 4 | 0.48 | 0.07 |
| Subthalamic nucleus | 4 | 0.59 | 0.58 |
| Suprachiasmatic nucleus | 2 | 0.61 | -2.32 |
| Supramammillary nucleus | 1 | 0.37 | 0.05 |
| Supraoptic nucleus | 4 | 0.45 | -2.10 |
| Tuberal nucleus | 4 | 0.49 | 0.64 |
| Ventrolateral preoptic nucleus | 1 | 0.22 | -2.82 |
| Ventromedial hypothalamic nucleus | 4 | 0.49 | 0.77 |
| Zona incerta | 4 | 0.50 | 0.85 |
| Anterior pretectal nucleus | 2 | 0.68 | -0.63 |
| Cuneiform nucleus | 4 | 0.40 | 0.44 |
| Inferior colliculus | 1 | 0.19 | 0.38 |
| Interpeduncular nucleus | 1 | 0.26 | 0.06 |
| Medial pretectal area | 1 | 0.33 | -1.59 |
| Midbrain reticular nucleus | 1 | 0.37 | 0.26 |
| Midbrain reticular nucleus retrorubral area | 1 | 0.26 | -0.19 |
| Nucleus of Darkschewitsch | 2 | 0.69 | -2.09 |
| Nucleus of the brachium of the inferior colliculus | 1 | 0.22 | 0.26 |
| Nucleus of the optic tract | 3 | 0.40 | 0.67 |
| Nucleus of the posterior commissure | 3 | 0.49 | 0.20 |
| Olivary pretectal nucleus | 3 | 0.44 | 0.33 |
| Parabigeminal nucleus | 3 | 0.38 | 0.20 |
| Pedunculopontine nucleus | 1 | 0.21 | 1.08 |
| Periaqueductal gray | 1 | 0.15 | 0.78 |
| Posterior pretectal nucleus | 3 | 0.66 | -2.16 |
| Precommissural nucleus | 3 | 0.46 | 1.21 |
| Red nucleus | 1 | 0.17 | 0.68 |
| Substantia nigra compact part | 2 | 0.64 | 0.19 |
| Substantia nigra reticular part | 4 | 0.62 | -0.15 |
| Superior colliculus motor related | 1 | 0.36 | 0.45 |
| Superior colliculus sensory related | 1 | 0.25 | 0.64 |
| Ventral tegmental area | 1 | 0.37 | -0.33 |
| Pons | 1 | 0.18 | 1.06 |
| Pons motor related | 1 | 0.26 | 0.75 |
| Pontine reticular nucleus | 1 | 0.23 | 0.82 |
| Vestibular nuclei | 1 | 0.47 | -1.51 |
| Ansiform lobule | 1 | 0.25 | 0.84 |
| Central lobule | 1 | 0.20 | 0.76 |
| Culmen | 1 | 0.19 | 0.40 |
| Paraflocculus | 1 | 0.27 | -0.03 |
| Simple lobule | 1 | 0.36 | -0.48 |
