## Supplemental Table 6 for "Characterization of the brain functional architecture of psychostimulant withdrawal using single-cell whole brain imaging"

**Supplementary Table 6: Top to bottom order of brain regions in Fig 1**

| **Number** | **Allen Order (Fig 1)** |
| --- | --- |
| 1 | Agranular insular area posterior part |
| 2 | Agranular insular area ventral part |
| 3 | Anterior cingulate area dorsal part |
| 4 | Anterior cingulate area ventral part |
| 5 | Anterior olfactory nucleus |
| 6 | Anterolateral visual area |
| 7 | Anteromedial visual area |
| 8 | Cortical amygdalar area posterior part |
| 9 | Dentate gyrus |
| 10 | Dorsal auditory area |
| 11 | Dorsal peduncular area |
| 12 | Ectorhinal area |
| 13 | Entorhinal area lateral part |
| 14 | Entorhinal area medial part |
| 15 | Fasciola cinerea |
| 16 | Field CA1 |
| 17 | Field CA2 |
| 18 | Field CA3 |
| 19 | Frontal pole cerebral cortex |
| 20 | Gustatory areas |
| 21 | Induseum griseum |
| 22 | Infralimbic area |
| 23 | Lateral visual area |
| 24 | Nucleus of the lateral olfactory tract |
| 25 | Orbital area lateral part |
| 26 | Orbital area medial part |
| 27 | Orbital area ventrolateral part |
| 28 | Parasubiculum |
| 29 | Perirhinal area |
| 30 | Piriform area |
| 31 | Piriform-amygdalar area |
| 32 | Posterior auditory area |
| 33 | Posterolateral visual area |
| 34 | Posteromedial visual area |
| 35 | Postpiriform transition area |
| 36 | Postsubiculum |
| 37 | Prelimbic area |
| 38 | Presubiculum |
| 39 | Primary auditory area |
| 40 | Primary motor area |
| 41 | Primary somatosensory area barrel field |
| 42 | Primary somatosensory area lower limb |
| 43 | Primary somatosensory area mouth |
| 44 | Primary somatosensory area nose |
| 45 | Primary somatosensory area trunk |
| 46 | Primary somatosensory area upper limb |
| 47 | Primary visual area |
| 48 | Retrosplenial area dorsal part |
| 49 | Retrosplenial area lateral agranular part |
| 50 | Retrosplenial area ventral part |
| 51 | Secondary motor area |
| 52 | Subiculum |
| 53 | Supplemental somatosensory area |
| 54 | Taenia tecta |
| 55 | Temporal association areas |
| 56 | Ventral auditory area |
| 57 | Visceral area |
| 58 | Basolateral amygdalar nucleus |
| 59 | Claustrum |
| 60 | Endopiriform nucleus |
| 61 | Lateral amygdalar nucleus |
| 62 | Posterior amygdalar nucleus |
| 63 | Anterior amygdalar area |
| 64 | Bed nucleus of the accessory olfactory tract |
| 65 | Caudoputamen |
| 66 | Central amygdalar nucleus |
| 67 | Fundus of striatum |
| 68 | Intercalated amygdalar nucleus |
| 69 | Lateral septal complex |
| 70 | Medial amygdalar nucleus |
| 71 | Nucleus accumbens |
| 72 | Olfactory tubercle |
| 73 | Septofimbrial nucleus |
| 74 | Bed nuclei of the stria terminalis |
| 75 | Diagonal band nucleus |
| 76 | Globus pallidus external segment |
| 77 | Globus pallidus internal segment |
| 78 | Magnocellular nucleus |
| 79 | Medial septal nucleus |
| 80 | Substantia innominata |
| 81 | Triangular nucleus of septum |
| 82 | Anterior group of the dorsal thalamus |
| 83 | Anterodorsal nucleus |
| 84 | Anteroventral nucleus of thalamus |
| 85 | Central lateral nucleus of the thalamus |
| 86 | Central medial nucleus of the thalamus |
| 87 | Dorsal part of the lateral geniculate complex |
| 88 | Interanterodorsal nucleus of the thalamus |
| 89 | Interanteromedial nucleus of the thalamus |
| 90 | Intergeniculate leaflet of the lateral geniculate complex |
| 91 | Intermediodorsal nucleus of the thalamus |
| 92 | Lateral dorsal nucleus of thalamus |
| 93 | Lateral habenula |
| 94 | Lateral posterior nucleus of the thalamus |
| 95 | Medial geniculate complex |
| 96 | Medial habenula |
| 97 | Mediodorsal nucleus of thalamus |
| 98 | Nucleus of reuniens |
| 99 | Paracentral nucleus |
| 100 | Parafascicular nucleus |
| 101 | Parataenial nucleus |
| 102 | Paraventricular nucleus of the thalamus |
| 103 | Peripeduncular nucleus |
| 104 | Posterior complex of the thalamus |
| 105 | Posterior limiting nucleus of the thalamus |
| 106 | Reticular nucleus of the thalamus |
| 107 | Submedial nucleus of the thalamus |
| 108 | Subparafascicular nucleus |
| 109 | Thalamus sensory-motor cortex related |
| 110 | Ventral anterior-lateral complex of the thalamus |
| 111 | Ventral medial nucleus of the thalamus |
| 112 | Ventral part of the lateral geniculate complex |
| 113 | Ventral posterior complex of the thalamus |
| 114 | Ventral posterolateral nucleus of the thalamus |
| 115 | Anterior hypothalamic nucleus |
| 116 | Anterodorsal preoptic nucleus |
| 117 | Anteroventral periventricular nucleus |
| 118 | Anteroventral preoptic nucleus |
| 119 | Arcuate hypothalamic nucleus |
| 120 | Dorsal premammillary nucleus |
| 121 | Dorsomedial nucleus of the hypothalamus |
| 122 | Lateral hypothalamic area |
| 123 | Lateral preoptic area |
| 124 | Mammillary body |
| 125 | Medial preoptic area |
| 126 | Medial preoptic nucleus |
| 127 | Median preoptic nucleus |
| 128 | Parastrial nucleus |
| 129 | Parasubthalamic nucleus |
| 130 | Paraventricular hypothalamic nucleus |
| 131 | Paraventricular hypothalamic nucleus descending division |
| 132 | Periventricular hypothalamic nucleus posterior part |
| 133 | Periventricular hypothalamic nucleus preoptic part |
| 134 | Periventricular zone |
| 135 | Posterior hypothalamic nucleus |
| 136 | Preparasubthalamic nucleus |
| 137 | Retrochiasmatic area |
| 138 | Subparaventricular zone |
| 139 | Subthalamic nucleus |
| 140 | Suprachiasmatic nucleus |
| 141 | Supramammillary nucleus |
| 142 | Supraoptic nucleus |
| 143 | Tuberal nucleus |
| 144 | Ventrolateral preoptic nucleus |
| 145 | Ventromedial hypothalamic nucleus |
| 146 | Zona incerta |
| 147 | Anterior pretectal nucleus |
| 148 | Cuneiform nucleus |
| 149 | Inferior colliculus |
| 150 | Interpeduncular nucleus |
| 151 | Medial pretectal area |
| 152 | Midbrain reticular nucleus |
| 153 | Midbrain reticular nucleus retrorubral area |
| 154 | Nucleus of Darkschewitsch |
| 155 | Nucleus of the brachium of the inferior colliculus |
| 156 | Nucleus of the optic tract |
| 157 | Nucleus of the posterior commissure |
| 158 | Olivary pretectal nucleus |
| 159 | Parabigeminal nucleus |
| 160 | Pedunculopontine nucleus |
| 161 | Periaqueductal gray |
| 162 | Posterior pretectal nucleus |
| 163 | Precommissural nucleus |
| 164 | Red nucleus |
| 165 | Substantia nigra compact part |
| 166 | Substantia nigra reticular part |
| 167 | Superior colliculus motor related |
| 168 | Superior colliculus sensory related |
| 169 | Ventral tegmental area |
| 170 | Pons |
| 171 | Pons motor related |
| 172 | Pontine reticular nucleus |
| 173 | Vestibular nuclei |
| 174 | Ansiform lobule |
| 175 | Central lobule |
| 176 | Culmen |
| 177 | Paraflocculus |
| 178 | Simple lobule |
