## Supplemental Table 7 for "Characterization of the brain functional architecture of psychostimulant withdrawal using single-cell whole brain imaging"

**Supplementary Table 7: Top to bottom order of brain regions in Fig 2**

| **Number** | **Saline Hierarchical Order** | **Cocaine Hierarchical Order** |
| --- | --- | --- |
| 1 | Retrosplenial area ventral part | Inferior colliculus |
| 2 | Interanterodorsal nucleus of the thalamus | Primary visual area |
| 3 | Anterior hypothalamic nucleus | Nucleus of the optic tract |
| 4 | Posterolateral visual area | Thalamus sensory-motor cortex related |
| 5 | Precommissural nucleus | Retrosplenial area dorsal part |
| 6 | Superior colliculus motor related | Field CA1 |
| 7 | Cuneiform nucleus | Retrosplenial area lateral agranular part |
| 8 | Primary visual area | Anterior amygdalar area |
| 9 | Superior colliculus sensory related | Anterodorsal preoptic nucleus |
| 10 | Parasubthalamic nucleus | Primary somatosensory area trunk |
| 11 | Vestibular nuclei | Interanteromedial nucleus of the thalamus |
| 12 | Pons motor related | Subparafascicular nucleus |
| 13 | Lateral visual area | Superior colliculus motor related |
| 14 | Anterolateral visual area | Periaqueductal gray |
| 15 | Pontine reticular nucleus | Magnocellular nucleus |
| 16 | Periaqueductal gray | Bed nuclei of the stria terminalis |
| 17 | Parastrial nucleus | Midbrain reticular nucleus retrorubral area |
| 18 | Fasciola cinerea | Ventromedial hypothalamic nucleus |
| 19 | Anterodorsal nucleus | Ventral tegmental area |
| 20 | Triangular nucleus of septum | Anterior pretectal nucleus |
| 21 | Lateral hypothalamic area | Endopiriform nucleus |
| 22 | Dorsomedial nucleus of the hypothalamus | Olfactory tubercle |
| 23 | Nucleus accumbens | Tuberal nucleus |
| 24 | Anterior group of the dorsal thalamus | Piriform area |
| 25 | Paraventricular hypothalamic nucleus descending division | Substantia innominata |
| 26 | Medial pretectal area | Ventral auditory area |
| 27 | Postsubiculum | Dorsal part of the lateral geniculate complex |
| 28 | Parasubiculum | Posterolateral visual area |
| 29 | Nucleus of the optic tract | Nucleus of the brachium of the inferior colliculus |
| 30 | Midbrain reticular nucleus retrorubral area | Supraoptic nucleus |
| 31 | Inferior colliculus | Cuneiform nucleus |
| 32 | Anterior pretectal nucleus | Paraventricular nucleus of the thalamus |
| 33 | Nucleus of Darkschewitsch | Lateral visual area |
| 34 | Field CA1 | Orbital area ventrolateral part |
| 35 | Nucleus of the posterior commissure | Red nucleus |
| 36 | Fundus of striatum | Parastrial nucleus |
| 37 | Dentate gyrus | Parasubthalamic nucleus |
| 38 | Presubiculum | Anterior hypothalamic nucleus |
| 39 | Lateral posterior nucleus of the thalamus | Posterior hypothalamic nucleus |
| 40 | Parafascicular nucleus | Dorsal premammillary nucleus |
| 41 | Peripeduncular nucleus | Lateral hypothalamic area |
| 42 | Central lobule | Retrochiasmatic area |
| 43 | Posterior pretectal nucleus | Perirhinal area |
| 44 | Lateral habenula | Field CA3 |
| 45 | Nucleus of reuniens | Posterior complex of the thalamus |
| 46 | Ventral medial nucleus of the thalamus | Entorhinal area lateral part |
| 47 | Tuberal nucleus | Intercalated amygdalar nucleus |
| 48 | Periventricular hypothalamic nucleus posterior part | Substantia nigra compact part |
| 49 | Posterior amygdalar nucleus | Basolateral amygdalar nucleus |
| 50 | Ventromedial hypothalamic nucleus | Pedunculopontine nucleus |
| 51 | Posterior hypothalamic nucleus | Medial preoptic area |
| 52 | Arcuate hypothalamic nucleus | Ectorhinal area |
| 53 | Subthalamic nucleus | Primary auditory area |
| 54 | Paracentral nucleus | Temporal association areas |
| 55 | Substantia nigra compact part | Pontine reticular nucleus |
| 56 | Culmen | Substantia nigra reticular part |
| 57 | Pedunculopontine nucleus | Pons |
| 58 | Interpeduncular nucleus | Midbrain reticular nucleus |
| 59 | Ventral posterior complex of the thalamus | Field CA2 |
| 60 | Induseum griseum | Supramammillary nucleus |
| 61 | Preparasubthalamic nucleus | Anteromedial visual area |
| 62 | Nucleus of the brachium of the inferior colliculus | Posterior auditory area |
| 63 | Red nucleus | Visceral area |
| 64 | Ventral tegmental area | Primary motor area |
| 65 | Substantia innominata | Paraventricular hypothalamic nucleus descending division |
| 66 | Medial geniculate complex | Lateral dorsal nucleus of thalamus |
| 67 | Subiculum | Primary somatosensory area barrel field |
| 68 | Midbrain reticular nucleus | Orbital area medial part |
| 69 | Thalamus sensory-motor cortex related | Orbital area lateral part |
| 70 | Simple lobule | Anterolateral visual area |
| 71 | Paraflocculus | Median preoptic nucleus |
| 72 | Submedial nucleus of the thalamus | Suprachiasmatic nucleus |
| 73 | Subparafascicular nucleus | Supplemental somatosensory area |
| 74 | Olivary pretectal nucleus | Agranular insular area posterior part |
| 75 | Central lateral nucleus of the thalamus | Primary somatosensory area lower limb |
| 76 | Medial septal nucleus | Septofimbrial nucleus |
| 77 | Subparaventricular zone | Anterior cingulate area ventral part |
| 78 | Anterior cingulate area ventral part | Paraventricular hypothalamic nucleus |
| 79 | Secondary motor area | Primary somatosensory area upper limb |
| 80 | Suprachiasmatic nucleus | Submedial nucleus of the thalamus |
| 81 | Periventricular zone | Nucleus accumbens |
| 82 | Septofimbrial nucleus | Claustrum |
| 83 | Paraventricular hypothalamic nucleus | Agranular insular area ventral part |
| 84 | Orbital area lateral part | Lateral septal complex |
| 85 | Mediodorsal nucleus of thalamus | Taenia tecta |
| 86 | Posteromedial visual area | Arcuate hypothalamic nucleus |
| 87 | Retrosplenial area dorsal part | Olivary pretectal nucleus |
| 88 | Anteroventral periventricular nucleus | Dorsomedial nucleus of the hypothalamus |
| 89 | Bed nuclei of the stria terminalis | Prelimbic area |
| 90 | Retrosplenial area lateral agranular part | Periventricular hypothalamic nucleus preoptic part |
| 91 | Medial preoptic nucleus | Gustatory areas |
| 92 | Anterodorsal preoptic nucleus | Frontal pole cerebral cortex |
| 93 | Primary motor area | Subparaventricular zone |
| 94 | Lateral septal complex | Caudoputamen |
| 95 | Primary somatosensory area lower limb | Fundus of striatum |
| 96 | Lateral dorsal nucleus of thalamus | Infralimbic area |
| 97 | Primary somatosensory area trunk | Medial septal nucleus |
| 98 | Anteromedial visual area | Central lateral nucleus of the thalamus |
| 99 | Lateral preoptic area | Posteromedial visual area |
| 100 | Periventricular hypothalamic nucleus preoptic part | Lateral posterior nucleus of the thalamus |
| 101 | Median preoptic nucleus | Central lobule |
| 102 | Infralimbic area | Central medial nucleus of the thalamus |
| 103 | Primary somatosensory area upper limb | Periventricular hypothalamic nucleus posterior part |
| 104 | Supramammillary nucleus | Cortical amygdalar area posterior part |
| 105 | Gustatory areas | Nucleus of the lateral olfactory tract |
| 106 | Taenia tecta | Entorhinal area medial part |
| 107 | Supraoptic nucleus | Zona incerta |
| 108 | Claustrum | Ventral anterior-lateral complex of the thalamus |
| 109 | Anteroventral nucleus of thalamus | Posterior amygdalar nucleus |
| 110 | Prelimbic area | Postpiriform transition area |
| 111 | Piriform area | Lateral preoptic area |
| 112 | Agranular insular area ventral part | Parabigeminal nucleus |
| 113 | Dorsal peduncular area | Intergeniculate leaflet of the lateral geniculate complex |
| 114 | Anterior cingulate area dorsal part | Ventral part of the lateral geniculate complex |
| 115 | Orbital area medial part | Interanterodorsal nucleus of the thalamus |
| 116 | Orbital area ventrolateral part | Lateral amygdalar nucleus |
| 117 | Anterior amygdalar area | Ventrolateral preoptic nucleus |
| 118 | Caudoputamen | Central amygdalar nucleus |
| 119 | Primary somatosensory area barrel field | Dorsal auditory area |
| 120 | Agranular insular area posterior part | Preparasubthalamic nucleus |
| 121 | Paraventricular nucleus of the thalamus | Ventral posterior complex of the thalamus |
| 122 | Medial habenula | Interpeduncular nucleus |
| 123 | Frontal pole cerebral cortex | Peripeduncular nucleus |
| 124 | Anterior olfactory nucleus | Dentate gyrus |
| 125 | Central medial nucleus of the thalamus | Superior colliculus sensory related |
| 126 | Intercalated amygdalar nucleus | Piriform-amygdalar area |
| 127 | Medial preoptic area | Medial geniculate complex |
| 128 | Intermediodorsal nucleus of the thalamus | Posterior pretectal nucleus |
| 129 | Supplemental somatosensory area | Nucleus of Darkschewitsch |
| 130 | Primary somatosensory area nose | Posterior limiting nucleus of the thalamus |
| 131 | Primary somatosensory area mouth | Paracentral nucleus |
| 132 | Visceral area | Subiculum |
| 133 | Dorsal auditory area | Ansiform lobule |
| 134 | Entorhinal area lateral part | Diagonal band nucleus |
| 135 | Field CA2 | Medial preoptic nucleus |
| 136 | Mammillary body | Paraflocculus |
| 137 | Posterior auditory area | Medial amygdalar nucleus |
| 138 | Ventral auditory area | Globus pallidus internal segment |
| 139 | Temporal association areas | Nucleus of reuniens |
| 140 | Ventral posterolateral nucleus of the thalamus | Mammillary body |
| 141 | Ansiform lobule | Globus pallidus external segment |
| 142 | Entorhinal area medial part | Reticular nucleus of the thalamus |
| 143 | Intergeniculate leaflet of the lateral geniculate complex | Presubiculum |
| 144 | Perirhinal area | Pons motor related |
| 145 | Reticular nucleus of the thalamus | Mediodorsal nucleus of thalamus |
| 146 | Ectorhinal area | Ventral medial nucleus of the thalamus |
| 147 | Posterior complex of the thalamus | Retrosplenial area ventral part |
| 148 | Ventral anterior-lateral complex of the thalamus | Nucleus of the posterior commissure |
| 149 | Dorsal part of the lateral geniculate complex | Parafascicular nucleus |
| 150 | Primary auditory area | Culmen |
| 151 | Postpiriform transition area | Simple lobule |
| 152 | Magnocellular nucleus | Precommissural nucleus |
| 153 | Globus pallidus internal segment | Vestibular nuclei |
| 154 | Lateral amygdalar nucleus | Parasubiculum |
| 155 | Nucleus of the lateral olfactory tract | Ventral posterolateral nucleus of the thalamus |
| 156 | Bed nucleus of the accessory olfactory tract | Bed nucleus of the accessory olfactory tract |
| 157 | Dorsal premammillary nucleus | Anteroventral preoptic nucleus |
| 158 | Substantia nigra reticular part | Subthalamic nucleus |
| 159 | Zona incerta | Anterior group of the dorsal thalamus |
| 160 | Ventral part of the lateral geniculate complex | Parataenial nucleus |
| 161 | Parabigeminal nucleus | Anteroventral periventricular nucleus |
| 162 | Field CA3 | Postsubiculum |
| 163 | Pons | Anterior cingulate area dorsal part |
| 164 | Retrochiasmatic area | Secondary motor area |
| 165 | Medial amygdalar nucleus | Triangular nucleus of septum |
| 166 | Parataenial nucleus | Primary somatosensory area mouth |
| 167 | Interanteromedial nucleus of the thalamus | Medial pretectal area |
| 168 | Piriform-amygdalar area | Anterior olfactory nucleus |
| 169 | Diagonal band nucleus | Primary somatosensory area nose |
| 170 | Ventrolateral preoptic nucleus | Anteroventral nucleus of thalamus |
| 171 | Anteroventral preoptic nucleus | Periventricular zone |
| 172 | Cortical amygdalar area posterior part | Intermediodorsal nucleus of the thalamus |
| 173 | Globus pallidus external segment | Medial habenula |
| 174 | Posterior limiting nucleus of the thalamus | Anterodorsal nucleus |
| 175 | Endopiriform nucleus | Fasciola cinerea |
| 176 | Olfactory tubercle | Dorsal peduncular area |
| 177 | Central amygdalar nucleus | Induseum griseum |
| 178 | Basolateral amygdalar nucleus | Lateral habenula |

| **Number** | **Methamphetamine Hierarchical Order** | **Nicotine Hierarchical Order** |
| --- | --- | --- |
| 1 | Caudoputamen | Ventral tegmental area |
| 2 | Anterior amygdalar area | Midbrain reticular nucleus retrorubral area |
| 3 | Parataenial nucleus | Superior colliculus motor related |
| 4 | Periventricular hypothalamic nucleus preoptic part | Midbrain reticular nucleus |
| 5 | Claustrum | Simple lobule |
| 6 | Medial habenula | Posterior hypothalamic nucleus |
| 7 | Medial pretectal area | Basolateral amygdalar nucleus |
| 8 | Ventral part of the lateral geniculate complex | Pedunculopontine nucleus |
| 9 | Anteroventral preoptic nucleus | Subparafascicular nucleus |
| 10 | Parasubthalamic nucleus | Pons motor related |
| 11 | Precommissural nucleus | Anterior cingulate area dorsal part |
| 12 | Parastrial nucleus | Paraventricular hypothalamic nucleus |
| 13 | Anteroventral periventricular nucleus | Ansiform lobule |
| 14 | Central amygdalar nucleus | Presubiculum |
| 15 | Lateral amygdalar nucleus | Dorsal auditory area |
| 16 | Endopiriform nucleus | Supplemental somatosensory area |
| 17 | Paraventricular nucleus of the thalamus | Posterior limiting nucleus of the thalamus |
| 18 | Intercalated amygdalar nucleus | Intercalated amygdalar nucleus |
| 19 | Intermediodorsal nucleus of the thalamus | Central amygdalar nucleus |
| 20 | Postpiriform transition area | Posteromedial visual area |
| 21 | Intergeniculate leaflet of the lateral geniculate complex | Lateral visual area |
| 22 | Ventral auditory area | Supramammillary nucleus |
| 23 | Bed nucleus of the accessory olfactory tract | Anterolateral visual area |
| 24 | Basolateral amygdalar nucleus | Gustatory areas |
| 25 | Dorsal auditory area | Mammillary body |
| 26 | Primary somatosensory area barrel field | Postsubiculum |
| 27 | Magnocellular nucleus | Periventricular hypothalamic nucleus posterior part |
| 28 | Primary somatosensory area nose | Periventricular zone |
| 29 | Induseum griseum | Paraflocculus |
| 30 | Anterior cingulate area ventral part | Peripeduncular nucleus |
| 31 | Anterior cingulate area dorsal part | Vestibular nuclei |
| 32 | Pedunculopontine nucleus | Anteroventral nucleus of thalamus |
| 33 | Superior colliculus motor related | Endopiriform nucleus |
| 34 | Inferior colliculus | Cortical amygdalar area posterior part |
| 35 | Entorhinal area lateral part | Postpiriform transition area |
| 36 | Substantia innominata | Prelimbic area |
| 37 | Nucleus accumbens | Intermediodorsal nucleus of the thalamus |
| 38 | Central lobule | Lateral septal complex |
| 39 | Posterior hypothalamic nucleus | Entorhinal area lateral part |
| 40 | Substantia nigra compact part | Ventrolateral preoptic nucleus |
| 41 | Parabigeminal nucleus | Visceral area |
| 42 | Parasubiculum | Posterior auditory area |
| 43 | Presubiculum | Temporal association areas |
| 44 | Postsubiculum | Primary auditory area |
| 45 | Diagonal band nucleus | Ventral auditory area |
| 46 | Posterior auditory area | Ectorhinal area |
| 47 | Piriform-amygdalar area | Perirhinal area |
| 48 | Periaqueductal gray | Pontine reticular nucleus |
| 49 | Supramammillary nucleus | Medial geniculate complex |
| 50 | Anterolateral visual area | Anterior cingulate area ventral part |
| 51 | Primary auditory area | Claustrum |
| 52 | Ectorhinal area | Pons |
| 53 | Medial geniculate complex | Central lobule |
| 54 | Temporal association areas | Red nucleus |
| 55 | Perirhinal area | Retrosplenial area lateral agranular part |
| 56 | Agranular insular area ventral part | Lateral amygdalar nucleus |
| 57 | Paraventricular hypothalamic nucleus | Retrosplenial area dorsal part |
| 58 | Subparafascicular nucleus | Interpeduncular nucleus |
| 59 | Subparaventricular zone | Superior colliculus sensory related |
| 60 | Paraventricular hypothalamic nucleus descending division | Inferior colliculus |
| 61 | Nucleus of the brachium of the inferior colliculus | Retrosplenial area ventral part |
| 62 | Midbrain reticular nucleus | Periaqueductal gray |
| 63 | Anterior hypothalamic nucleus | Ventral part of the lateral geniculate complex |
| 64 | Peripeduncular nucleus | Nucleus of the brachium of the inferior colliculus |
| 65 | Subiculum | Mediodorsal nucleus of thalamus |
| 66 | Lateral visual area | Culmen |
| 67 | Superior colliculus sensory related | Fasciola cinerea |
| 68 | Midbrain reticular nucleus retrorubral area | Agranular insular area posterior part |
| 69 | Nucleus of reuniens | Piriform area |
| 70 | Zona incerta | Central medial nucleus of the thalamus |
| 71 | Culmen | Interanteromedial nucleus of the thalamus |
| 72 | Retrosplenial area lateral agranular part | Medial pretectal area |
| 73 | Lateral preoptic area | Thalamus sensory-motor cortex related |
| 74 | Anterior pretectal nucleus | Septofimbrial nucleus |
| 75 | Posterior limiting nucleus of the thalamus | Ventral posterolateral nucleus of the thalamus |
| 76 | Preparasubthalamic nucleus | Piriform-amygdalar area |
| 77 | Nucleus of the optic tract | Dorsomedial nucleus of the hypothalamus |
| 78 | Medial preoptic area | Dentate gyrus |
| 79 | Thalamus sensory-motor cortex related | Anteromedial visual area |
| 80 | Medial preoptic nucleus | Posterolateral visual area |
| 81 | Dorsomedial nucleus of the hypothalamus | Fundus of striatum |
| 82 | Red nucleus | Caudoputamen |
| 83 | Lateral septal complex | Arcuate hypothalamic nucleus |
| 84 | Central medial nucleus of the thalamus | Parasubthalamic nucleus |
| 85 | Interpeduncular nucleus | Suprachiasmatic nucleus |
| 86 | Reticular nucleus of the thalamus | Subiculum |
| 87 | Medial septal nucleus | Medial septal nucleus |
| 88 | Supraoptic nucleus | Nucleus of reuniens |
| 89 | Periventricular hypothalamic nucleus posterior part | Substantia nigra compact part |
| 90 | Interanteromedial nucleus of the thalamus | Dorsal premammillary nucleus |
| 91 | Secondary motor area | Paraventricular hypothalamic nucleus descending division |
| 92 | Field CA2 | Central lateral nucleus of the thalamus |
| 93 | Field CA3 | Nucleus of Darkschewitsch |
| 94 | Posteromedial visual area | Anterior pretectal nucleus |
| 95 | Primary motor area | Parafascicular nucleus |
| 96 | Anteromedial visual area | Intergeniculate leaflet of the lateral geniculate complex |
| 97 | Medial amygdalar nucleus | Precommissural nucleus |
| 98 | Piriform area | Lateral habenula |
| 99 | Posterior amygdalar nucleus | Medial habenula |
| 100 | Primary somatosensory area trunk | Parabigeminal nucleus |
| 101 | Nucleus of the lateral olfactory tract | Nucleus of the optic tract |
| 102 | Primary somatosensory area upper limb | Nucleus of the posterior commissure |
| 103 | Primary somatosensory area lower limb | Olivary pretectal nucleus |
| 104 | Cortical amygdalar area posterior part | Anterodorsal nucleus |
| 105 | Visceral area | Posterior pretectal nucleus |
| 106 | Agranular insular area posterior part | Parataenial nucleus |
| 107 | Gustatory areas | Induseum griseum |
| 108 | Supplemental somatosensory area | Triangular nucleus of septum |
| 109 | Primary somatosensory area mouth | Paraventricular nucleus of the thalamus |
| 110 | Anterior olfactory nucleus | Interanterodorsal nucleus of the thalamus |
| 111 | Interanterodorsal nucleus of the thalamus | Medial preoptic area |
| 112 | Globus pallidus external segment | Lateral preoptic area |
| 113 | Anterodorsal preoptic nucleus | Nucleus accumbens |
| 114 | Mediodorsal nucleus of thalamus | Ventral medial nucleus of the thalamus |
| 115 | Ventral posterolateral nucleus of the thalamus | Globus pallidus internal segment |
| 116 | Median preoptic nucleus | Lateral hypothalamic area |
| 117 | Orbital area medial part | Anteroventral periventricular nucleus |
| 118 | Infralimbic area | Magnocellular nucleus |
| 119 | Prelimbic area | Dorsal peduncular area |
| 120 | Taenia tecta | Primary motor area |
| 121 | Fundus of striatum | Primary somatosensory area upper limb |
| 122 | Lateral habenula | Nucleus of the lateral olfactory tract |
| 123 | Olivary pretectal nucleus | Median preoptic nucleus |
| 124 | Entorhinal area medial part | Anterodorsal preoptic nucleus |
| 125 | Periventricular zone | Primary somatosensory area lower limb |
| 126 | Pons | Zona incerta |
| 127 | Dorsal premammillary nucleus | Agranular insular area ventral part |
| 128 | Pontine reticular nucleus | Field CA3 |
| 129 | Substantia nigra reticular part | Ventromedial hypothalamic nucleus |
| 130 | Lateral hypothalamic area | Parastrial nucleus |
| 131 | Ventral tegmental area | Primary visual area |
| 132 | Dentate gyrus | Taenia tecta |
| 133 | Lateral posterior nucleus of the thalamus | Field CA1 |
| 134 | Subthalamic nucleus | Field CA2 |
| 135 | Suprachiasmatic nucleus | Anteroventral preoptic nucleus |
| 136 | Posterolateral visual area | Retrochiasmatic area |
| 137 | Pons motor related | Infralimbic area |
| 138 | Ventromedial hypothalamic nucleus | Anterior amygdalar area |
| 139 | Retrochiasmatic area | Primary somatosensory area nose |
| 140 | Primary visual area | Submedial nucleus of the thalamus |
| 141 | Olfactory tubercle | Primary somatosensory area mouth |
| 142 | Retrosplenial area dorsal part | Secondary motor area |
| 143 | Field CA1 | Subparaventricular zone |
| 144 | Mammillary body | Primary somatosensory area trunk |
| 145 | Globus pallidus internal segment | Reticular nucleus of the thalamus |
| 146 | Arcuate hypothalamic nucleus | Periventricular hypothalamic nucleus preoptic part |
| 147 | Ventrolateral preoptic nucleus | Preparasubthalamic nucleus |
| 148 | Cuneiform nucleus | Anterior group of the dorsal thalamus |
| 149 | Tuberal nucleus | Posterior amygdalar nucleus |
| 150 | Submedial nucleus of the thalamus | Tuberal nucleus |
| 151 | Dorsal part of the lateral geniculate complex | Paracentral nucleus |
| 152 | Retrosplenial area ventral part | Cuneiform nucleus |
| 153 | Paraflocculus | Subthalamic nucleus |
| 154 | Bed nuclei of the stria terminalis | Substantia nigra reticular part |
| 155 | Anteroventral nucleus of thalamus | Entorhinal area medial part |
| 156 | Simple lobule | Parasubiculum |
| 157 | Fasciola cinerea | Orbital area medial part |
| 158 | Dorsal peduncular area | Globus pallidus external segment |
| 159 | Triangular nucleus of septum | Olfactory tubercle |
| 160 | Orbital area ventrolateral part | Supraoptic nucleus |
| 161 | Posterior pretectal nucleus | Dorsal part of the lateral geniculate complex |
| 162 | Nucleus of the posterior commissure | Medial preoptic nucleus |
| 163 | Nucleus of Darkschewitsch | Posterior complex of the thalamus |
| 164 | Frontal pole cerebral cortex | Orbital area lateral part |
| 165 | Anterior group of the dorsal thalamus | Ventral anterior-lateral complex of the thalamus |
| 166 | Vestibular nuclei | Orbital area ventrolateral part |
| 167 | Ventral posterior complex of the thalamus | Lateral dorsal nucleus of thalamus |
| 168 | Orbital area lateral part | Substantia innominata |
| 169 | Ansiform lobule | Diagonal band nucleus |
| 170 | Ventral anterior-lateral complex of the thalamus | Anterior olfactory nucleus |
| 171 | Anterodorsal nucleus | Primary somatosensory area barrel field |
| 172 | Septofimbrial nucleus | Anterior hypothalamic nucleus |
| 173 | Paracentral nucleus | Medial amygdalar nucleus |
| 174 | Posterior complex of the thalamus | Bed nuclei of the stria terminalis |
| 175 | Ventral medial nucleus of the thalamus | Ventral posterior complex of the thalamus |
| 176 | Central lateral nucleus of the thalamus | Frontal pole cerebral cortex |
| 177 | Lateral dorsal nucleus of thalamus | Lateral posterior nucleus of the thalamus |
| 178 | Parafascicular nucleus | Bed nucleus of the accessory olfactory tract |
